# The SC-SNc pathway boosts appetitive locomotion in predatory hunting

**DOI:** 10.1101/2020.11.23.395004

**Authors:** Meizhu Huang, Dapeng Li, Qing Pei, Zhiyong Xie, Huating Gu, Aixue Liu, Zijun Chen, Yi Wang, Fangmiao Sun, Yulong Li, Jiayi Zhang, Miao He, Yuan Xie, Fan Zhang, Xiangbing Qi, Congping Shang, Peng Cao

## Abstract

Appetitive locomotion is essential for organisms to approach rewards, such as food and prey. How the brain controls appetitive locomotion is poorly understood. In a naturalistic goal-directed behavior—predatory hunting, we demonstrate an excitatory brain circuit from the superior colliculus (SC) to the substantia nigra pars compacta (SNc) to boost appetitive locomotion. The SC-SNc pathway transmitted locomotion-speed signals to dopamine neurons and triggered dopamine release in the dorsal striatum. Activation of this pathway increased the speed and frequency of approach during predatory hunting, an effect that depended on the activities of SNc dopamine neurons. Conversely, synaptic inactivation of this pathway impaired appetitive locomotion but not defensive or exploratory locomotion. Together, these data revealed the SC as an important source to provide locomotion-related signals to SNc dopamine neurons to boost appetitive locomotion.

## INTRODUCTION

Locomotion plays a fundamental role in the survival of organisms. It can be conceptually divided into three categories: appetitive locomotion, defensive locomotion, and exploratory locomotion (Sinnamon, 1993). These three types of locomotion may be selectively recruited by distinct brain circuits for specific behavioral needs (Ferreira-Pinto et al., 2018). Appetitive locomotion is indispensible for organisms to approach rewarding targets. For example, in a naturalistic goal-directed behavior—predatory hunting, predator employs appetitive locomotion to chase and catch up with prey (Hoy et al., 2016; Han et al., 2017). How the brain control appetitive locomotion during predatory hunting is an unresolved question in the field of neuroethology (Sillar et al., 2016).

The superior colliculus (SC) is a multi-layered midbrain structure for sensory information processing and motor functions (Cang et al., 2018; Gandhi and Katnani, 2011; Basso and May, 2017). The superficial layers of the SC primarily receive visual inputs (Morin and Studholme, 2014) and perform visual information processing (Wang et al., 2010; De Franceschi and Solomon, 2018). The intermediate and deep layers of the SC are involved in sensorimotor transformation and motor functions (Gandhi and Katnani 2011). The motor functions of the SC include saccadic eye movement (Wurtz and Albano, 1980; Sparks, 1986; Wang et al., 2015), head movement (Isa and Sasaki, 2002; Wilson et al., 2018) and locomotion (Cooper et al., 1998; Felsen and Mainen, 2008). From a neuroethological perspective, these motor functions of the SC enable itself to orchestrate distinct behavioral actions in predatory hunting in rodents (Furigo et al., 2010; Favaro et al., 2011; Hoy et al., 2019; Shang et al., 2019).

How the SC orchestrates distinct behavioral actions during predatory hunting (e.g. approaching and attacking prey) is beginning to be elucidated. With an unbiased activity-dependent genetic labeling approach (FosTRAP2), several hunting-associated tectofugal pathways were identified, such as those projecting to the zona incerta (ZI) and the substantia nigra pars compacta (SNc) (Shang et al., 2019). While the SC-ZI pathway is primarily involved in sensory-triggered predatory attack during hunting, the functional role of the SC-SNc pathway in predatory hunting has not been determined yet.

The SC-SNc pathway, also known as tectonigral pathway, was first described by Comoli et al. (2003). It was shown that neurons in the intermediate and deep layers of the SC form synaptic contacts with dopamine and non-dopamine SNc neurons (Comoli et al., 2003). Considering the recent studies showing the involvement of SNc dopamine neurons in the vigor of body movements, including locomotion (Jin and Costa, 2010; Dodson et al., 2016; Howe and Dombeck, 2016; da Silva et al., 2018; Coddington and Dudman, 2018), we hypothesized that the SC-SNc pathway may participate in appetitive locomotion during predatory hunting.

In the present study, we explored the role of SC-SNc pathway in appetitive locomotion during predatory hunting. We found that the SC-SNc pathway transmitted locomotion speed signals to SNc dopamine neurons and triggered dopamine release in the dorsal striatum. Activation of this pathway during predatory hunting increased the speed of appetitive locomotion, an effect that depended on the activities of SNc dopamine neurons. Conversely, synaptic inactivation of this pathway impaired appetitive locomotion without changing defensive locomotion. Together, these data revealed the SC as an important source to provide locomotion-related signals to SNc dopamine neurons to boost appetitive locomotion.

## RESULTS

### The SC-SNc pathway is primarily glutamatergic

We began this study by performing morphological analyses of the SC-SNc pathway. First, we mapped the SC-SNc pathway with cell-type-specific expression of “SynaptoTag” (Xu and Südhof, 2013), which is the enhanced green fluorescent protein fused to synaptic vesicle protein synaptobrevin-2 (EGFP-Syb2). AAV-DIO-EGFP-Syb2 was unilaterally injected into the SC of *vGlut2-IRES-Cre* or *GAD2-IRES-Cre* mice (Figure 1A and 1C). The specificities of *vGlut2-IRES-Cre* and *GAD2-IRES-Cre* mice to label glutamate+ and GABA+ SC neurons have been validated in an earlier study (Shang et al., 2019). EGFP-Syb2 expression in SC neurons of *vGlut2-IRES-Cre* mice resulted in considerable EGFP-Syb2+ puncta in the SNc (Figure 1B and S1A). In contrast, only sparse EGFP-Syb2+ puncta were observed in the SNc of *GAD2-IRES-Cre* mice (Figure 1D and S1B). We normalized the density of EGFP-Syb2 puncta by dividing the puncta density in the SNc with that in the SC of each mouse. Strikingly, the normalized density of EGFP-Syb2 puncta in the SNc of *vGlut2-IRES-Cre* mice was significantly higher than that of *GAD2-IRES-Cre* mice, suggesting that the SC-SNc pathway is primarily glutamatergic (Figure 1E).

**Figure 1.**
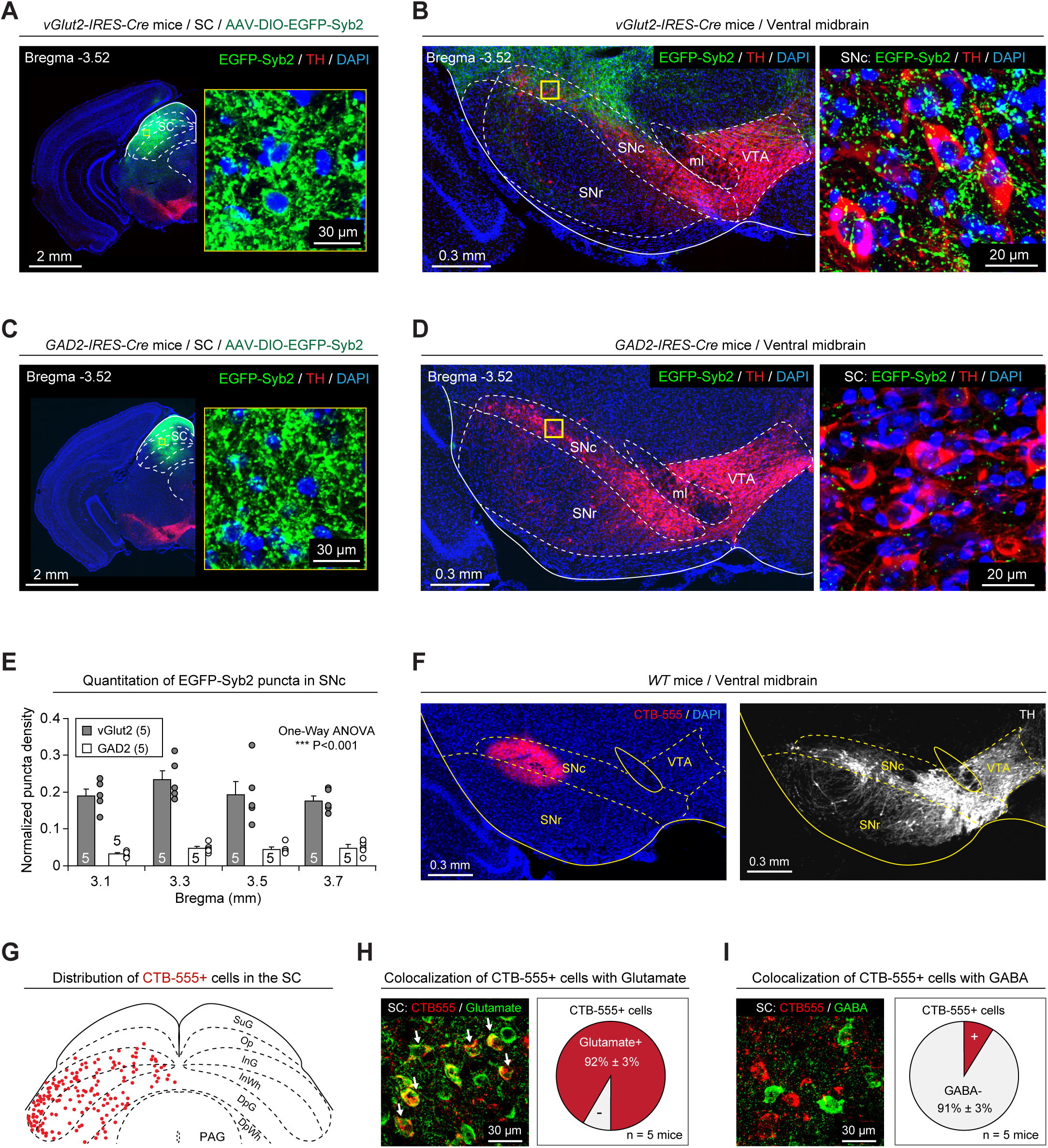
Cell-type-specific mapping of SC-SNc pathway. **(A, C)** Example coronal brain sections of *vGlut2-IRES-Cre* (A) and *GAD2-IRES-Cre* mice (C) with EGFP-Syb2 expression in the SC. Insets, high-magnification micrographs showing EGFP-Syb2+ puncta in the SC. **(B, D)** *Left*, EGFP-Syb2+ axon terminals in the ventral midbrain of *vGlut2-IRES-Cre* (B) or *GAD2-IRES-Cre* mice (D). *Right*, high-magnification micrographs showing EGFP-Syb2+ puncta (green) in the SNc. The boundary of the SNc was delineated according to the immunofluorescence of TH (red). **(E)** Normalized density of EGFP-Syb2+ puncta in the SNc of *vGlut2-IRES-Cre* (vGlut2) and *GAD2-IRES-Cre* (GAD2) mice as a function of bregma. The normalization was made by dividing the puncta density in the SNc with that in the SC. (**F**) An example coronal section of ventral midbrain showing the injection of CTB-555 into the SNc of *WT* mice (*left*). The boundaries of SNc and VTA were determined by immunofluorescence of TH of dopamine neurons (*right*). (**G**) A schematic diagram showing the distribution of SNc-projecting SC neurons that were labeled by CTB-555. For the raw image corresponding to the schematic diagram, see Figure S1C. (**H, I**) Example micrographs (*left*) and quantitative analyses (*right*) showing CTB-555+ cells in the SC are predominantly positive for glutamate (H) and negative for GABA (I). Number of mice was indicated in the graphs (E, H, I). Data in (E, H, I) are means ± SEM (error bars). Statistical analyses in (E) were performed by One-Way ANOVA (*** P < 0.001). For the P values, see Table S4. Scale bars are indicated in the graphs.

Second, we retrogradely labeled SNc-projecting SC neurons by injecting CTB-555 into the SNc of *WT* mice (Figure 1F). The retrogradely labeled cells (CTB-555+) in the SC were distributed predominantly in the intermediate and deep layers (Figure 1G and S1C). By using primary antibodies that specifically recognize GABA and glutamate (Shang et al., 2018), we found that most of the CTB-555+ cells were immunohistochemically glutamate+ (92% ± 3%, n = 5 mice; Figure 1H) and GABA− (91% ± 3%, n = 5 mice; Figure 1I). These data, again, suggested that the SC-SNc pathway is primarily glutamatergic.

In addition to the SC-SNc pathway, some SC neurons form another pathway to the ventral tegmental area (VTA) that was implicated in the regulation of sleep and innate defensive responses (Zhang et al., 2019; Zhou et al., 2019). To test whether SNc-projecting SC neurons send collaterals to the VTA, we injected CTB-488 and CTB-555 into the SNc and VTA of the same *WT* mice (Figure S2A). Interestingly, very few cells were dually labeled by CTB-555 and CTB-488 in the SC (Figure S2, B-E). As a negative control, injection of mixed CTB-555 and CTB-488 in the SNc (Figure S2F) resulted in SC neurons co-labeled by both CTB-555 and CTB-488 (Figure S2, G-J). These data suggested that the SC-SNc pathway is anatomically segregated from the SC-VTA pathway.

We also examined whether the SNc-projecting SC neurons send collaterals to the ZI, an important center for feeding-related predation (Zhang et al., 2017; Zhao et al., 2019). CTB-488 and CTB-555 were injected into the SNc and ZI of the same mice (Figure S3A). Again, very few cells in the SC were co-labeled by CTB-555 and CTB-488 (Figure S3, B-D), suggesting that the SNc-projecting SC neurons rarely send collaterals to the ZI.

### Single SNc-projecting SC neurons encode locomotion speed

To test whether the SC-SNc pathway is involved in locomotion, we made single-unit recording from the SNc-projecting SC neurons by using antidromic activation strategy (Shang et al., 2019). AAV-ChR2-mCherry (Boyden et al., 2005) was injected into the SC of *WT* mice, followed by implantation of an optical fiber above the ChR2-mCherry+ axon terminals in the SNc (Figure S4A). Three weeks after viral injection, single-unit recording was performed with a tungsten electrode in the SC of head-fixed awake mice walking on a cylindrical treadmill (Figure 2A, *left*). The putative SNc-projecting SC neurons were identified by the antidromic action potentials (APs) evoked by light pulses (473 nm, 1 ms, 2 mW) that illuminated ChR2-mCherry+ axon terminals in the SNc (Figure 2A, *right*). The antidromically evoked APs had to conform to two criteria (Cohen et al., 2012; Roseberry et al., 2016): First, their waveform should be similar to that of APs during locomotion; second, their latencies to light pulses should be less than 5 ms. With these empirical criteria, we identified 18 units as putative SNc-projecting SC neurons. Their antidromically evoked APs possessed waveforms quantitatively correlated with those of APs during locomotion (Figure S4B and 2B) and had short response latencies to light pulses (2.7 ms ± 0.4 ms, n=18 units; Figure 2C).

**Figure 2.**
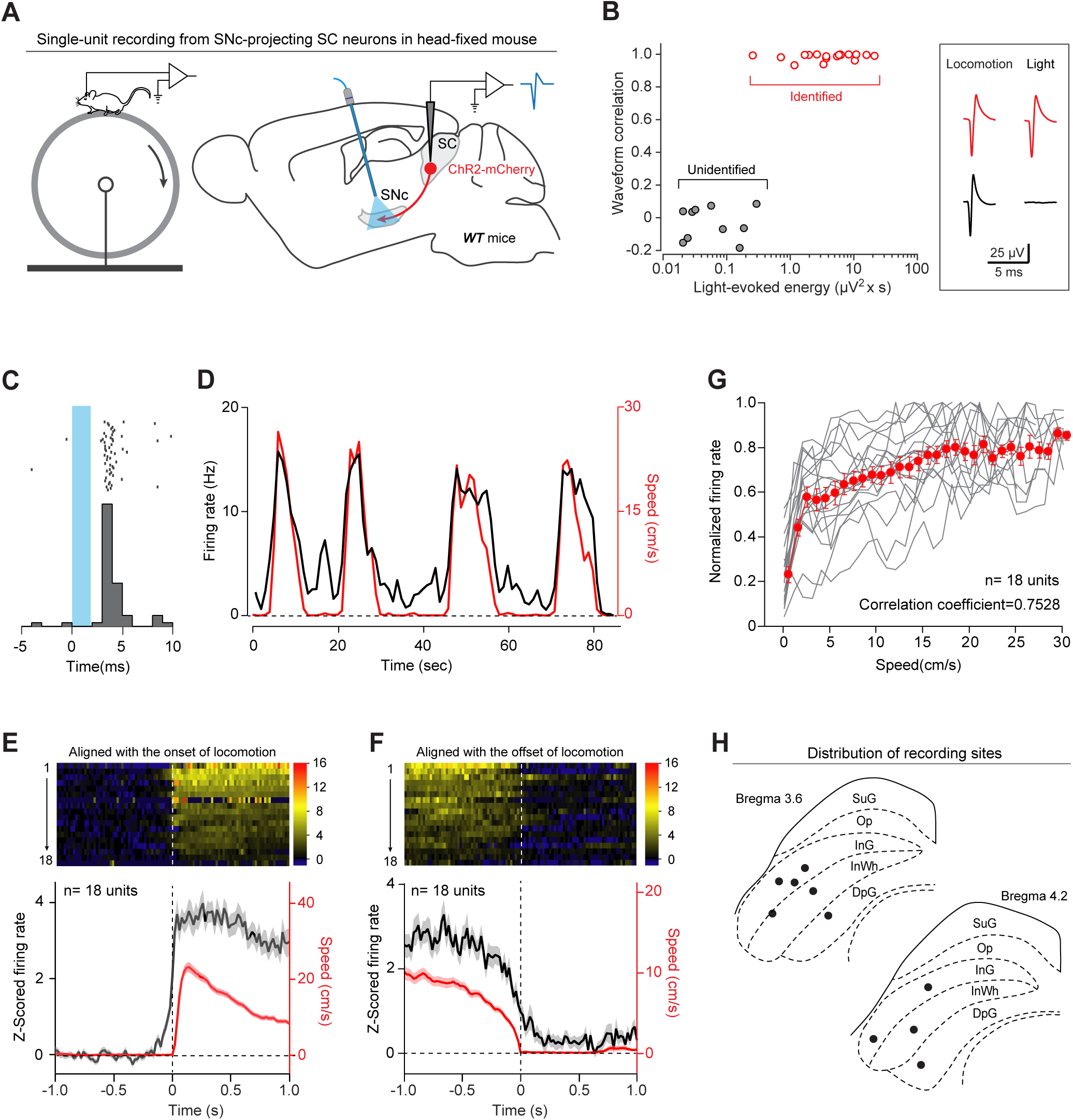
SNc-projecting SC neurons encode locomotion speed. **(A)** Schematic diagram showing a head-fixed awake mouse walking on a cylindrical treadmill (*left*) and antidromic activation strategy for single-unit recording from SNc-projecting SC neurons (*right*). **(B)** Correlation analysis of action potentials of individual units evoked either by light pulses (Light) or by locomotion (Locomotion), confirming a segregation between antidromically identified units (red dots and traces) and unidentified units (grey dots and traces). **(C)** Raster (*top*) and peristimulus time histogram (PSTH, *bottom*) of an example putative SNc-projecting SC neurons showing a latency of ∼3 ms relative to the onset of light pulses. **(D)** Smoothed PSTH (trace in black) of an example putative SNc-projecting SC neuron aligned with locomotion speed (trace in red). **(E, F)** Heat-map graphs (*top*) and averaged time course (*bottom*) of Z-scored firing rate of 18 putative SNc-projecting SC neurons aligned to the onset (E) and the offset (F) of locomotion. **(G)** Averaged (red) and individual (gray) normalized instantaneous firing rate during locomotion as a function of locomotion speed in 500 ms bins. **(H)** Distribution of recording site (black dots) in the SC. For example micrograph, see Figure S4C. Number of units was indicated in the graphs (E-G). Data in (E-G) are means ± SEM (error bars).

Then we examined the instantaneous firing rate of these putative SNc-projecting SC neurons before, during and after locomotion on the treadmill. In general, the activities of SNc-projecting SC neurons was modulated by locomotion (Movie S1; Figure 2D). To examine the temporal relationship between the activities of SNc-projecting SC neurons and locomotion initiation, we aligned firing rate of individual units with the onset of locomotion (Figure 2E, *top*). We defined the response onset time as the time when the signal reached 15% of peak amplitude relative to the baseline. The average response curve started to rise at 107 ms ± 15 ms before locomotion onset (n = 18 units; Figure 2E, *bottom*). Similarly, we aligned firing rate of individual units with the offset of locomotion (Figure 2F, *top*), and found that the average response curve dropped to baseline at 121 ms ± 16 ms after locomotion offset (n = 18 units; Figure 2F, *bottom*). To examine how the activities of these units were modulated during locomotion, we plotted the response-speed curve of each single-unit (Caggiano et al., 2018) and found a correlation between the firing rate and locomotion speed in the range of 3 cm/s ∼ 30 cm/s (Figure 2G; Spearman correlation coefficient=0.7528; P=7.06E-07). Histological verification indicated that all the recorded units were localized within the intermediate and deep layers of the SC (Figure 2H and S4C). These data suggested that the SNc-projecting SC neurons encode locomotion speed of mice.

### The SC-SNc pathway promotes appetitive locomotion in predatory hunting

To further explore the role of SC-SNc pathway in locomotion, we examined whether activation of this pathway promotes locomotion. AAV-ChR2-mCherry was bilaterally injected into the SC of *WT* mice (Figure 3A and S5A), followed by implantation of an optical fiber above the ChR2-mCherry+ axon terminals in the SNc (Figure 3B). In acute brain slices with the SC, light pulses (10 Hz, 2 ms, 10 mW) reliably evoked action potentials from ChR2-mCherry+ SC neurons (Figure S5B). In a linear runway (Figure 3C), photostimulation of the SC-SNc pathway (10 Hz, 20 ms, 6 s, 10 mW) increased locomotion speed of mice (Figure 3D and 3E). In control mice injected with AAV-mCherry in the SC, light illumination on mCherry+ axon terminals in the SNc did not promote locomotion speed (Figure 3D and 3E). In addition, we found that the effects of SC-SNc pathway activation on locomotion speed depended on the frequency of light stimulation (Figure S5C).

**Figure 3.**
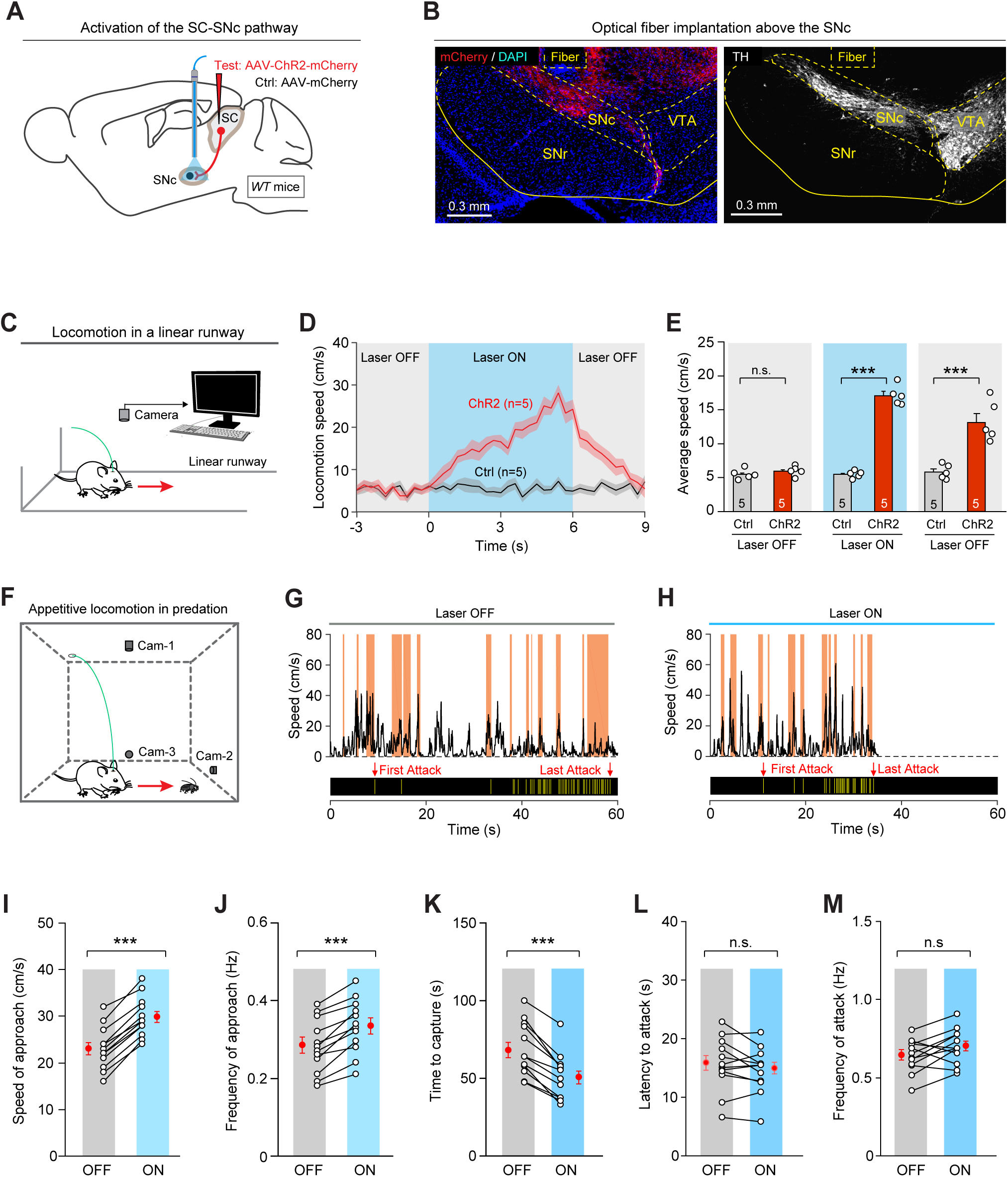
Activation of the SC-SNc pathway promoted appetitive locomotion. **(A)** Schematic diagram showing injection of AAV-ChR2-mCherry into the SC of *WT* mice, followed by optical fiber implantation above the SNc. For the example micrograph of ChR2-mCherry expression in the SC, see Figure S5A. (**B**) An example coronal section of ventral midbrain with an optical-fiber track above the ChR2-mCherry+ axon terminals in the SNc (*left*), the boundary of which was delineated by the immunofluorescence of TH (*right*). (**C**) Schematic diagram showing the experimental configuration to monitor mouse locomotor behavior in the linear runway. **(D)** Time courses of locomotion speed of control (Ctrl) and test mice (ChR2) in the linear runway before, during and after light stimulation of the SC-SNc pathway (10 Hz, 10 ms, 6 s, 10 mW). (**E**) Quantitative analyses of average locomotion speed of control (Ctrl) and test mice (ChR2) before, during and after photostimulation of the SC-SNc pathway. For the dependence of locomotion speed on the frequency of photostimulation, see Figure S5C. (**F**) Schematic diagram showing the experimental configuration to monitor predatory hunting in the arena. (**G, H**) Time course of locomotion speed (*top*) and jaw attacks (*bottom*) during predatory hunting of an example mouse without (G) and with (H) photostimulation of the SC-SNc pathway (10 Hz, 10 ms, 10 mW). The shaded areas (orange) indicated the approach episodes in predatory hunting. For the analyses of azimuth angle and PPD, see Figure S5F and S5G. (**I-M**) Speed of approach (I), frequency of approach (J), time to capture (K), latency to attack (L), and frequency of attack (M) in predatory hunting of mice without (OFF) and with (ON) photostimulation of the SC-SNc pathway. Number of mice was indicated in the graphs (D, E, I-M). Data in (D, E, I-M) are means ± SEM (error bars). Statistical analyses in (E, I-M) were performed by Student t-tests (n.s. P>0.1; *** P < 0.001). For the P values, see Table S4. Scale bars are indicated in the graphs.

Then we tested whether activation of the SC-SNc pathway boosts appetitive locomotion during predatory hunting. In predatory hunting, appetitive locomotion occurred when predator approached prey (Figure 3F). By measuring the instantaneous azimuth angle and distance between prey and predator (Figure S5D), we were able to identify a series of intermittent approach episodes (Figure S5E) according to the established criteria (Hoy et al., 2016). Appetitive locomotion in these approach episodes was quantitatively assessed by measuring the speed of approach and frequency of approach. The speed of approach was calculated by averaging the peak speed of each approach episode in the trial. The frequency of approach was the number of approach episodes divided by total time of the trial. With the method above, we labeled the approach episodes (shaded areas in orange) in the behavioral ethogram of predatory hunting in mice without (OFF) and with (ON) photostimulation of the SC-SNc pathway (10 Hz, 20 ms, 10 mW) (Movie S2 and S3; Figure 3G and 3H). We found that activation of the SC-SNc pathway significantly increased the speed of approach (Figure 3I), increased the frequency of approach (Figure 3J) and reduced the time required for prey capture (Figure 3K). In contrast, the latency and the frequency of predatory attack with jaw during hunting were not altered by activation of the SC-SNc pathway (Figure 3, L and M). These data suggested that activation of the SC-SNc pathway specifically promoted appetitive locomotion during predatory hunting.

### The SC-SNc pathway is required for appetitive locomotion in predatory hunting

Then we examined whether the SC-SNc pathway is required for appetitive locomotion during predatory hunting, by synaptically inactivating the SNc-projecting SC neurons. Tetanus neurotoxin (TeNT), which blocks neurotransmitter release by proteolytic cleavage of synaptobrevin-2 (Schiavo et al., 1992), has been used as a molecular tool for synaptic inactivation (Xu and Südhof, 2013; Cregg et al., 2020). The effectiveness and specificity of TeNT-mediated synaptic inactivation of SC neurons have been validated in a previous study (Shang et al., 2019). To synaptically inactivate the SC-SNc pathway, we injected AAV2-retro-mCherry-IRES-Cre (Tervo et al, 2016) and AAV-DIO-EGFP-2A-TeNT into the SNc and SC of *WT* mice, respectively (Figure 4A). The injection of AAV2-retro-mCherry-IRES-Cre in the SNc (Figure S6A) and AAV-DIO-EGFP-2A-TeNT in the SC cooperatively labeled SNc-projecting SC neurons with EGFP (Figure 4B, *top*), as demonstrated by the co-expression of mCherry and EGFP in the same SC neurons (Figure 4B, *bottom*).

**Figure 4.**
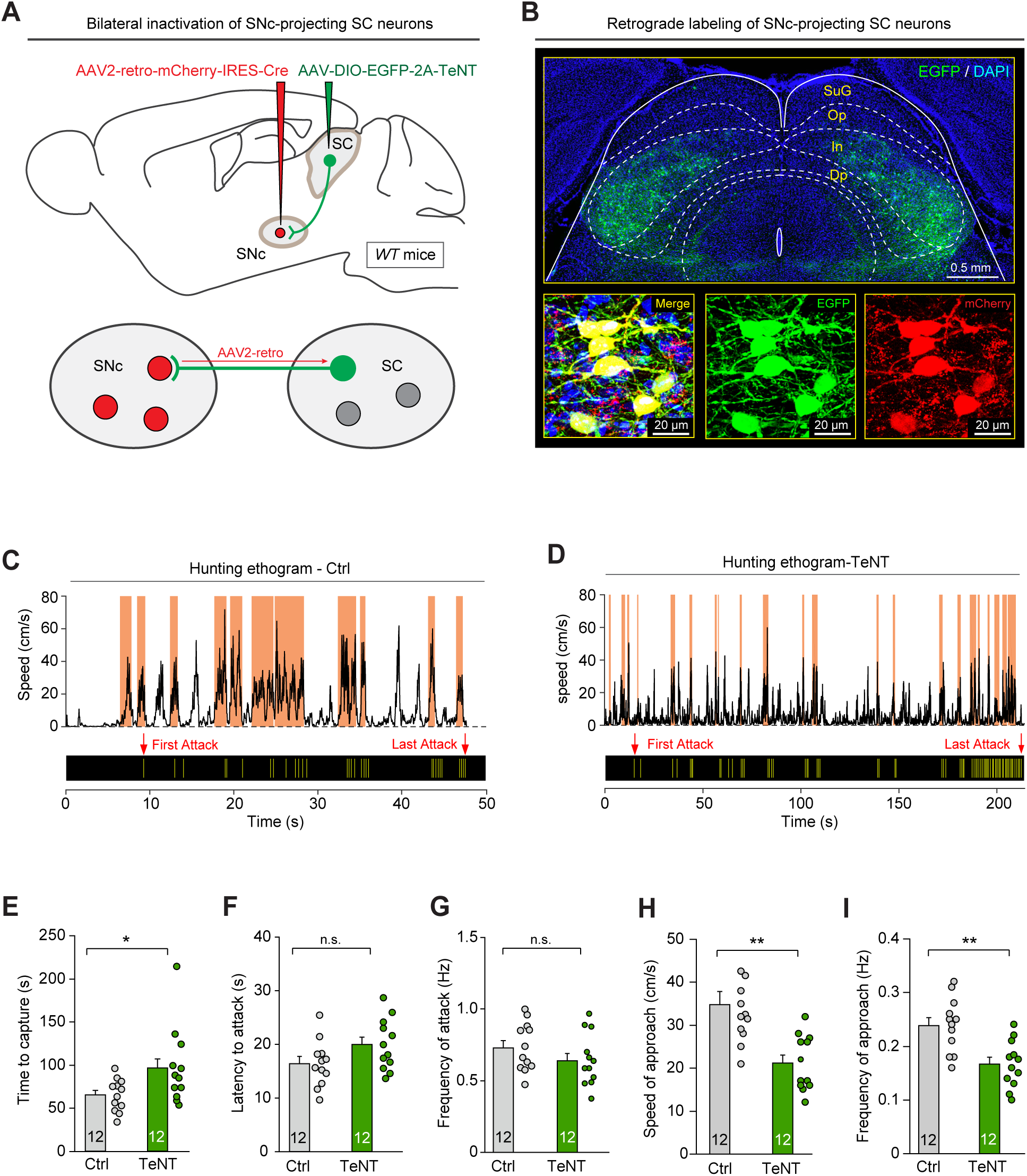
The SC-SNc pathway is required for appetitive locomotion during predatory hunting. **(A)** Schematic diagram showing the AAV2-retro strategy to selectively inactivate the SNc-projecting SC neurons with TeNT. For the example coronal section of ventral midbrain showing the injection of AAV2-retro-mCherry-IRES-Cre, see Figure S6A. (**B**) An example coronal brain section showing EGFP+ SNc-projecting SC neurons distributed in the intermediate layer (In) and deep layer (Dp) of SC. Inset, merged and single-channel micrographs showing SNc-projecting SC neurons were dually labeled by mCherry and EGFP. **(C, D)** Time course of locomotion speed (*top*) and jaw attacks (*bottom*) during predatory hunting of example mice either without (C, Ctrl) or with (D, TeNT) synaptic inactivation of SNc-projecting SC neurons. The shaded areas (orange) indicated the approach episodes in predatory hunting. For the analyses of azimuth angle and PPD, see Figure S6B and S6C. (**E-I**) Quantitative analyses of time to capture (E), latency to attack (F), frequency of attack (G), speed of approach (H), and frequency of approach (I) during predatory hunting in mice without (Ctrl) and with (TeNT) synaptic inactivation of SNc-projecting SC neurons. Number of mice was indicated in the graphs (E-I). Data in (E-I) are means ± SEM (error bars). Statistical analyses in (E-I) were performed by Student t-tests (n.s. P>0.1; * P<0.05; ** P < 0.01). For the P values, see Table S4. Scale bars are indicated in the graphs.

Then we analyzed the effects of SC-SNc pathway inactivation on predatory hunting behavior in mice. We labeled the approach episodes (shaded areas in orange) in the behavioral ethogram of predatory hunting in mice without (Ctrl) and with (TeNT) synaptic inactivation of the SC-SNc pathway (Movie S4 and S5; Figure 4C and 4D). TeNT-mediated inactivation of SC-SNc pathway impaired predatory hunting by significantly increasing the time required for prey capture (Figure 4E). Such effect on hunting could not be explained by an impairment of predatory attack, because neither the latency nor the frequency of jaw attack during hunting was changed by SC-SNc pathway inactivation (Figure 4F and 4G). In contrast, both the speed of approach and frequency of approach in predatory hunting were significantly decreased (Figure 4H and 4I). These data suggested that the SC-SNc pathway is required for appetitive locomotion during predatory hunting.

We also tested the role of the SC-SNc pathway in defensive and exploratory locomotion. In response to looming visual stimuli, mice exhibited escape behavior to avoid the imminent threats (Yilmaz and Meister, 2013). The escape behavior was used to measure defensive locomotion (Caggiano et al., 2018). We found that mice without (Ctrl) and with (TeNT) synaptic inactivation of the SC-SNc pathway exhibited escape followed by freezing in response to looming visual stimuli (Movie S6 and S7; Figure S6D and S6E). Quantitative analyses of locomotion speed indicated that synaptic inactivation of the SC-SNc pathway did not alter the peak speed of escape behavior during looming visual stimuli (Figure S6F) or average speed after stimuli (Figure S6G), suggesting this pathway is not required for defensive locomotion. Moreover, these mice exhibited similar average locomotion speed before looming visual stimuli (Figure S6H), suggesting that the SC-SNc pathway may not be involved in exploratory locomotion as well.

### The SC-SNc pathway preferentially innervates SNc dopamine neurons

To explore how the SC-SNc pathway is synaptically connected to the SNc, we employed mouse lines to genetically label different neuronal subtypes in the SNc. The most prominent neuronal subtype in the SNc is the dopamine neurons positive for tyrosine hydroxylase (TH+). These SNc dopamine neurons are largely segregated from those expressing glutamate decarboxylase-2 (GAD2+) (Tritsch et al., 2014; Kim et al., 2015) or vesicular glutamate transporter-2 (vGlut2+) (Kawano et al., 2006; Morales and Root, 2014). Although *TH-GFP* mice (Sawamoto et al., 2001) did not reliably label dopamine neurons in the VTA, this line marked SNc dopamine neurons with higher fidelity (Lammel et al., 2015). We confirmed this observation (Figure S7A and S7B) and used *TH-GFP* mice to genetically label SNc dopamine neurons in this study. By crossing *Ai14* (Madisen et al., 2010) with *GAD2-IRES-Cre* mice (Taniguchi et al., 2011), we labeled putative SNc GAD2+ neurons with tdTomato (GAD2-tdT; Figure S7C and S7D). Similarly, *Ai14* was crossed with *vGlut2-IRES-Cre* mice (Vong et al., 2011) to label putative SNc vGlut2+ neurons with tdTomato (vGlut2-tdT; Figure S7E and S7F).

To examine how the SC-SNc pathway synaptically innervates SNc dopamine neurons and GAD2+ neurons, we generated *GAD2-IRES-Cre/Ai14/TH-GFP* triple transgenic mice. In this mouse line, putative SNc dopamine neurons were genetically labeled by GFP (TH-GFP+), whereas putative GAD2+ neurons were identified as those positive for tdTomato (GAD2-tdT+) (Figure 5A). AAV-ChR2-mCherry was injected into the SC of *GAD2-IRES-Cre/Ai14/TH-GFP* mice (Figure 5C, *left*). In acute brain slice with the SNc, we illuminated ChR2-mCherry+ axon terminals with light-pulses (473 nm, 2 ms) with saturating power (20 mW), while performing whole-cell recordings from TH-GFP+ and adjacent GAD2-tdT+ neurons (Figure 5C, *right*). Using low-chloride internal solution (Kim et al., 2015), we recorded optogenetically-evoked excitatory postsynaptic currents (oEPSCs, voltage clamp at −70 mV) and inhibitory postsynaptic currents (oIPSCs, voltage clamp at 0 mV), which were removed by perfusion of antagonists of glutamate receptors (APV & CNQX) and GABAa receptor (picrotoxin), respectively (Figure S7, G-I). We found that the amplitude of oEPSCs was significantly higher than that of oIPSCs in both SNc GAD2-tdT+ and TH-GFP+ neurons (Figure 5, D and E). This was consistent with the morphological observation that the SNc-projecting SC neurons are primarily glutamatergic (Figure 1). Moreover, the amplitude of oEPSCs in TH-GFP+ neurons was significantly higher than that in GAD2-tdT+ neurons (Figure 5, D and E), suggesting that the SC-SNc pathway preferentially innervate SNc dopamine neurons.

**Figure 5.**
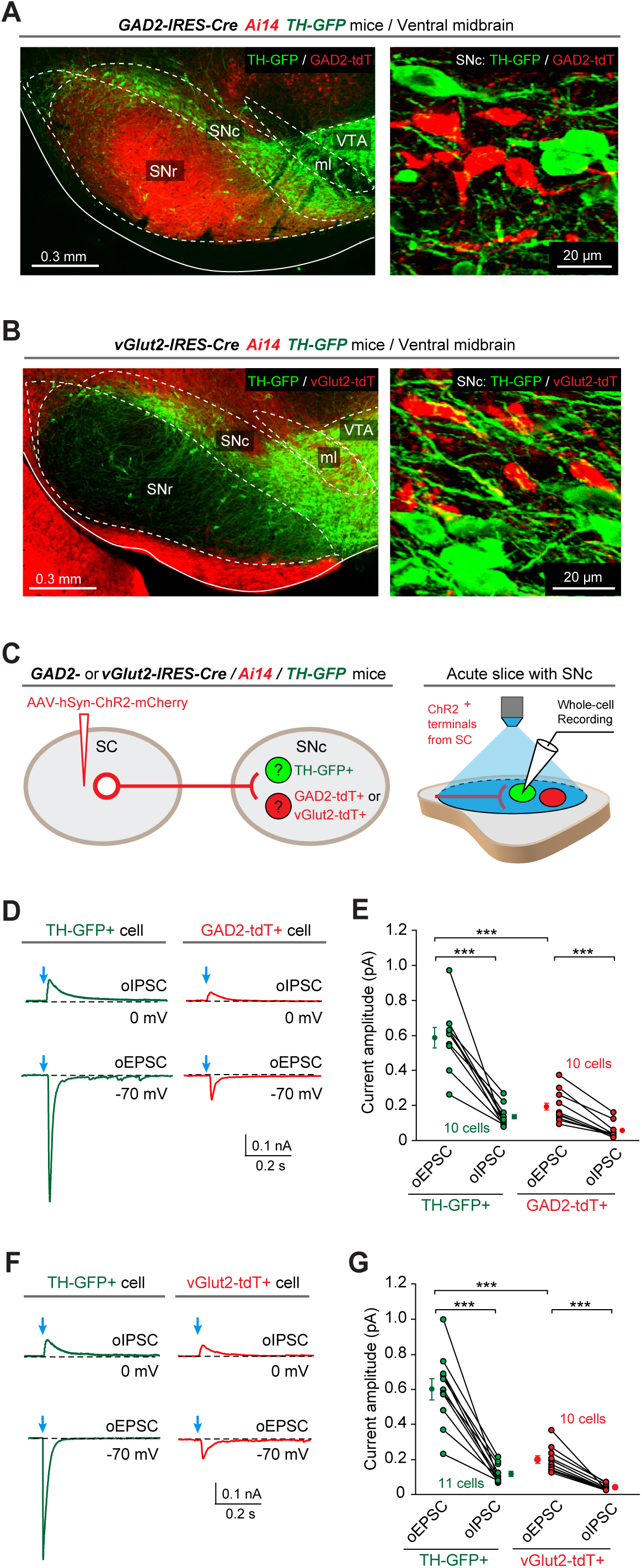
Dopamine neurons are the primary postsynaptic target of the SC-SNc pathway. (**A**) An example coronal section with ventral midbrain (*left*) and the high-magnification micrograph (*right*) showing the segregation of GAD2-tdT+ neurons and TH-GFP+ neurons in the SNc of *GAD2-IRES-Cre/Ai14/TH-GFP* triple transgenic mice. (**B**) An example coronal section with ventral midbrain (*left*) and the high-magnification micrograph (*right*) showing the segregation of vGlut2-tdT+ neurons and TH-GFP+ neurons in the SNc of *vGlut2-IRES-Cre/Ai14/TH-GFP* triple transgenic mice. **(C)** Schematic diagram showing injection of AAV-hSyn-ChR2-mCherry into the SC of *GAD2-IRES-Cre/Ai14/TH-GFP* or *vGlut2-IRES-Cre/Ai14/TH-GFP* mice (*left*) and whole-cell recording from TH-GFP+ (green), GAD2-tdT+ (red) or vGlut2-tdT+ (red) neurons while illuminating ChR2-positive axon terminals from the SC (*right*). **(D, E)** Example traces (D) and quantitative analyses (E) of oEPSCs and oIPSCs from TH-GFP+ and GAD2-tdT+ neurons in the slices of *GAD2-IRES-Cre/Ai14/TH-GFP* mice. **(F, G)** Example traces (F) and quantitative analyses (E) of oEPSCs and oIPSCs from TH-GFP+ and vGlut2-tdT+ neurons in the slices of *vGlut2-IRES-Cre/Ai14/TH-GFP* mice. Number of neurons is indicated in the graphs (E, G). Data in (E, G) are means ± SEM (error bars). Statistical analyses in (E, G) were performed by Student t-test (*** P < 0.001). For the P values, see Table S4. Scale bars are indicated in the graphs.

To test how the SC-SNc pathway is synaptically connected to SNc vGlut2+ neurons, we generated v*Glut2-IRES-Cre/Ai14/TH-GFP* triple transgenic mice. In this mouse line, putative SNc dopamine neurons were genetically labeled by GFP (TH-GFP+), whereas putative vGlut2+ neurons were identified as those positive for tdTomato (vGlut2-tdT+) (Figure 5B). We injected AAV-ChR2-mCherry into the SC of triple transgenic mice (Figure 5C, *left*). In acute brain slices with the SNc, we recorded oEPSCs and oIPSCs from TH-GFP+ and adjacent vGlut2-tdT+ neurons (Figure 5C, *right*). We found that the amplitude of oEPSCs was significantly higher than that of oIPSCs in both TH-GFP+ and vGlut2-tdT+ neurons (Figure 5, F and G). Moreover, the amplitude of oEPSCs in TH-GFP+ neurons was significantly higher than that in vGlut2-tdT+ neurons (Figure 5, F and G). These data indicated that the SC-SNc pathway has a stronger synaptic connection with SNc dopamine neurons than their synaptic connections to GAD2+ or vGlut2+ non-dopamine neurons.

### Activation of the SC-SNc pathway triggers striatal dopamine release

To further confirm that the SNc dopamine neurons are the postsynaptic target of SC-SNc pathway, we examined whether activation of this pathway evoke dopamine release in the dorsal striatum (Lerner et al., 2015). To monitor dopamine release, we employed genetically encoded GPCR-activation-based dopamine sensor (GRAB_DA_ sensor) that reports dopamine dynamics of nigrostriatal pathway (Sun et al., 2018). AAV-C1V1-mCherry (Yizhar et al., 2011) and AAV-GRAB_DA_ were injected into the SC and dorsal striatum of *WT* mice (Figure 6A), followed by implantation of optical fibers above the SNc and dorsal striatum, respectively (Figure 6B). The viral expression and optical fiber implantation were validated by using immunohistochemistry and slice physiology (Figure 6, C-F).

**Figure 6.**
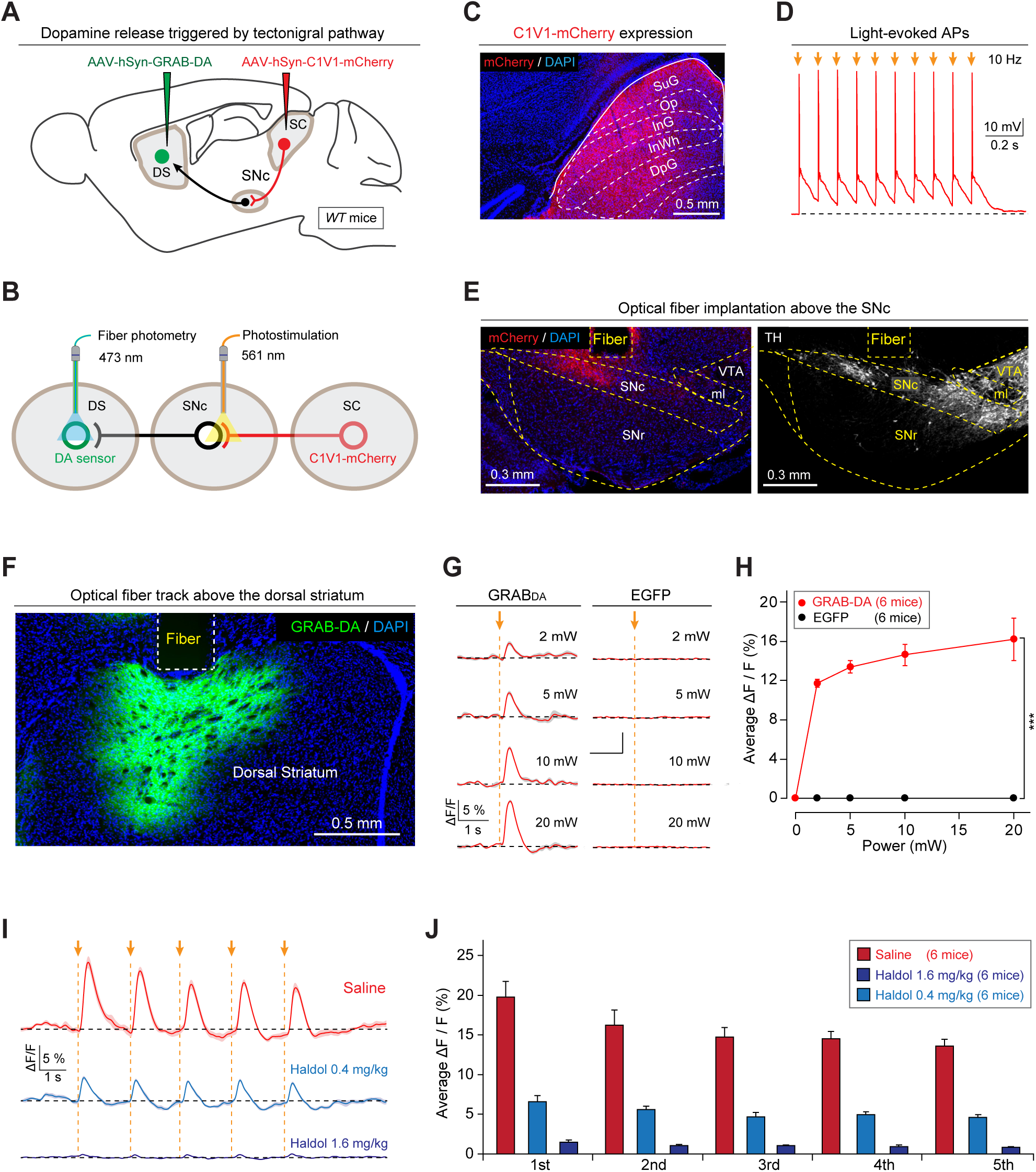
Activation of the SC-SNc pathway triggers dopamine release in the dorsal striatum. **(A)** Schematic diagram showing injections of AAV-hSyn-C1V1-mCherry in the SC and AAV-hSyn-GRAB-DA in the dorsal striatum (DS) of *WT* mice. **(B)** Schematic diagram showing implantation of optical fibers above the SNc and DS to apply photostimulation and fiber photometry recording, respectively. (**C**) An example coronal section with C1V1-mCherry expression in the SC. (**D**) In acute SC slices, light pulses (2 ms, 561 nm, 10 Hz, 10 pulses) illuminating on C1V1-mCherry+ SC neurons reliably triggered action potential firing. (**E**) An example coronal section of ventral midbrain showing an optical-fiber track above the C1V1-mCherry+ axon terminals in the SNc (*left*), the boundary of which was determined according to the immunofluorescence of TH (*right*). (**F**) An example coronal section with an optical-fiber track above the DS neurons expressing GRAB-DA sensor. **(G, H)** Example traces (G) and input-output curve (H) of GRAB-DA signals evoked by photostimulation of the SC-SNc pathway with different laser power. EGFP was used as a control of GRAB-DA sensor. **(I, J)** Example traces (I) and quantitative analyses (J) of GRAB-DA signals evoked by photostimulation (561 nm, 5 pulses, 2 ms, 0.5 Hz, 10 mW) of the SC-SNc pathway in mice treated with saline or Haloperidol (0.4 or 1.6 mg/kg). Number of mice was indicated in the graphs (H, J). Data in (H, J) are means ± SEM (error bars). Statistical analyses in (H, J) were performed by One-Way ANOVA (*** P < 0.001). For the P values, see Table S4. Scale bars are indicated in the graphs.

Then we tested whether activation of the SC-SNc pathway triggers dopamine release in the dorsal striatum. In freely moving mice, single light-pulses (561nm, 2 ms, 0∼20 mW) stimulating the axon terminals of SNc-projecting SC neurons (Figure 6, B and E) transiently increased the fluorescence of GRAB_DA_ sensor in the dorsal striatum (Figure 6, G and H). As a control, no obvious fluorescence changes were observed in striatal neurons expressing EGFP (Figure 6, G and H). Moreover, the light-evoked GRAB_DA_ signals were abrogated by D2 receptor antagonist haloperidol (Figure 6, I and J). These data indicated that SC-SNc pathway activation triggers dopamine release in the dorsal striatum, supporting that the SNc dopamine neurons are the postsynaptic target of the SC-SNc pathway.

### SNc dopamine neurons mediate appetitive locomotion evoked by SC-SNc pathway

Then we asked whether the SNc dopamine neurons mediate appetitive locomotion evoked by SC-SNc pathway. To address this question, we employed the strategy of designer receptors exclusively activated by designer drugs (DREADD) to chemogenetically silence SNc dopamine neurons (Armbruster et al., 2007). AAV-DIO-hM4Di-mCherry and AAV-ChR2-EYFP were injected into the SNc and SC of *DAT-IRES-Cre* mice (Backman et al., 2006) bilaterally, followed by two optical fibers implanted above the SNc (Figure 7, A and B). AAV-DIO-mCherry was used as a control of AAV-DIO-hM4Di-mCherry. In the SC, the expression of ChR2-EYFP and the efficiency to evoke action potentials from ChR2-EYFP+ neurons were validated (Figure S8A and S8B). In the ventral midbrain, hM4Di-mCherry was specifically expressed in SNc dopamine neurons that were intermingled with ChR2-EYFP+ axon terminals from SC neurons (Figure 7C). Chemogenetic suppression of neuronal firing by Clozapine N-oxide (CNO, 10 μM) was confirmed in slice physiology (Figure 7D).

**Figure 7.**
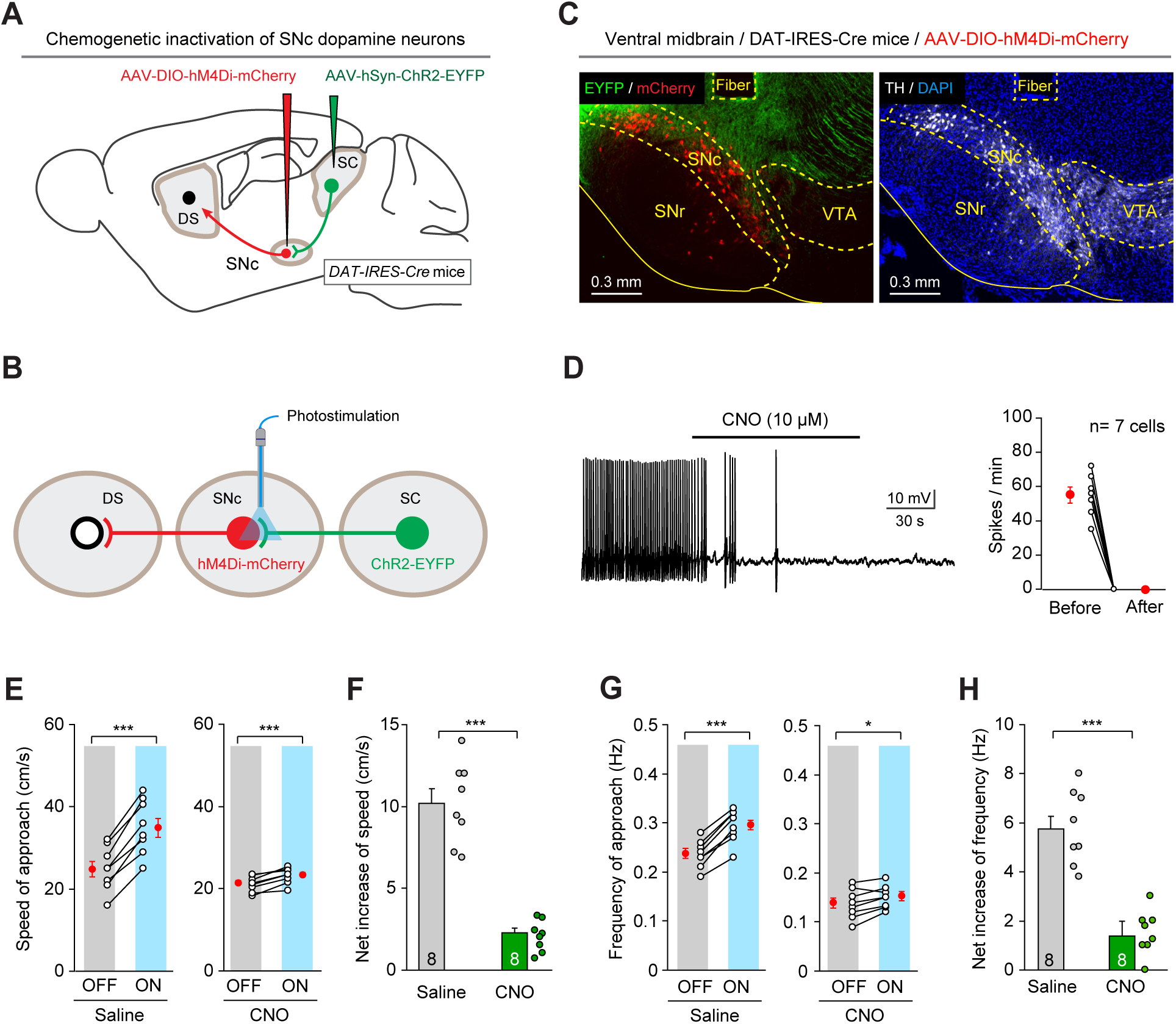
SNc dopamine neurons mediated the effect of SC-SNc pathway activation. **(A)** Schematic diagram showing injection of AAV-DIO-hM4Di-mCherry and AAV-ChR2-EYFP into the SNc and SC of *DAT-IRES-Cre* mice. For an example coronal section showing the expression of ChR2-EYFP in the SC, see Figure S8A. **(B)** Schematic diagram showing implantation of an optical fiber above the SNc to apply light stimulation on ChR2-EYFP+ axon terminals from the SC. (**C**) An example coronal section of ventral midbrain showing the optical fiber track above the hM4Di-mCherry+ SNc dopamine neurons intermingled with ChR2-EYFP+ axons from the SC (*left*). The boundaries of SNc and VTA were delineated by the immunofluorescence of TH (*right*). (**D**) An example train of action potentials recorded from SNc dopamine neurons expressing hM4Di-mCherry (*left*) and quantitative analyses of spike number per minute before and 3 min after perfusion of 10 μM CNO in ACSF (*right*). (**E**) The effect of light stimulation of the SC-SNc pathway on speed of approach during predatory hunting in mice treated with saline (*left*) or CNO (*right*). **(F)** Chemogenetic suppression of SNc dopamine neurons with CNO significantly attenuated the net effects of light stimulation of the SC-SNc pathway on speed of approach during predatory hunting. **(G)** The effect of light stimulation of the SC-SNc pathway on frequency of approach during predatory hunting in mice treated with saline (*left*) or CNO (*right*). **(H)** Chemogenetic suppression of SNc dopamine neurons with CNO significantly attenuated the net effects of light stimulation of the SC-SNc pathway on frequency of approach during predatory hunting. Number of cells (D) and mice (E-H) was indicated in the graphs. Data in (D-H) are means ± SEM (error bars). Statistical analyses in (D-H) were performed by Student t-tests (n.s. P>0.1; * P<0.05; *** P < 0.001). For the P values, see Table S4. Scale bars are indicated in the graphs.

To test whether SNc dopamine neurons mediate the appetitive locomotion evoked by SC-SNc pathway activation, we intraperitoneally treated the mice with saline or CNO. Light stimulation of SC-SNc pathway of mice treated with saline significantly increased the speed of approach (Figure 7E, *left*) and the frequency of approach (Figure 7G, *left*) during predatory hunting. When the same mice were treated with CNO (1 mg/kg) to chemogenetically suppress the activities of SNc dopamine neurons, activation of SC-SNc pathway only mildly increased the speed of approach (Figure 7E, *right*) and the frequency of approach (Figure 7G, *right*). For each mouse, we calculated “net increase” of approach speed by subtracting speed of approach during laser OFF from that during laser ON. It turned out that chemogenetic inactivation of SNc dopamine neurons with CNO prevented the net increase of approach speed (Figure 7F). Similarly, we calculated “net increase” of approach frequency by subtracting frequency of approach during laser OFF from that during laser ON. We found that inactivation of SNc dopamine neurons prevented the net increase of approach frequency during predatory hunting (Figure 7H). These data suggested that the activities of SNc dopamine neurons are required for the appetitive locomotion evoked by SC-SNc pathway.

## DISCUSSION

Appetitive locomotion is required for organisms to approach incentive stimuli in goal-directed behaviors. How the brain controls appetitive locomotion is poorly understood. Here we used predatory hunting as a behavior paradigm to address this question. We demonstrate an excitatory subcortical circuit from the SC to the SNc to boost appetitive locomotion. The SC-SNc pathway transmits locomotion-speed signals to dopamine neurons and triggers dopamine release in the dorsal striatum. Activation of this pathway promoted appetitive locomotion during predatory hunting, whereas synaptic inactivation of this pathway impairs appetitive locomotion rather than defensive locomotion. Together, these data reveal the SC as an important source to provide locomotion-related signals to SNc dopamine neurons to boost appetitive locomotion.

### The brain circuits for predatory hunting: the SC and beyond

As a naturalistic goal-directed behavior, predatory hunting has been the focus of studies using diverse animal models, such as toad (Ewert, 1997), zebrafish (Gahtan et al., 2005; Bianco et al., 2011; Trivedi and Bollmann, 2013) and rodents (Anjum et al., 2006; Hoy et al., 2016). In these animal models, it was found that the optic tectum (OT) and its mammalian homolog, the SC, play a fundamental role in predatory hunting (Toad: Ewert, 1997; Zebrafish: Del Bene et al., 2010; Bianco and Engert, 2015; Rodents: Furigo et al., 2010; Favaro et al., 2011). In rodents, a recent study has shown that genetically-defined neuronal subtypes in the SC make distinct contributions to prey capture behavior in mice (Hoy et al., 2019). The hunting-associated SC neurons may form divergent neural pathways to orchestrate distinct behavioral actions during predatory hunting, such as attacking prey (Shang et al., 2019) and, as demonstrated in this study, appetitive locomotion for approaching prey.

In another line of research, it was found that brain areas, which were thought to be related to food intake, are also involved in predatory hunting. For example, optogenetic activation of GABAergic neurons in the central amygdala (CeA), the lateral hypothalamus (LH), or the ZI provoked strong predatory hunting in mice (Han et al., 2017; Li et al., 2018; Zhao et al., 2019). The involvement of feeding-related areas in predatory hunting may be evolutionarily conserved, because the inferior lobe of hypothalamus in zebrafish also participates in prey capture behavior (Muto et al., 2017). In addition, activation of CaMKIIα-positive neurons in the medial preoptic area (MPA), which is related to object craving, also induces hunting-like actions toward prey (Park et al., 2018). Understanding how the neurons in the SC and in these newly-discovered brain areas coordinately control predatory hunting is a challenging task for future study.

### Dopamine system modulates predatory hunting

As an important neuromodulatory system in the brain, dopamine system plays a critical role in conditioned and unconditioned appetitive behaviors (Schultz, 2007; Bromberg-Martin et al., 2010). Earlier studies using systemic treatment of agonists or antagonists of dopamine receptors have demonstrated strong effects of dopaminergic modulation on predatory hunting in mammals (Schmidt, 1983; Shaikh et al., 1991; Tinsley et al., 2000). However, two critical questions remained unanswered. First, how is dopamine system recruited during predatory hunting? Second, considering the multiple clusters of dopamine neurons in the brain, which specific clusters of dopamine neurons participate in modulating predatory hunting? In this study, we show that the dopamine neurons in the SNc are innervated by the SC, a central hub to orchestrate predatory hunting. The SC-SNc pathway may provide locomotion-related signals to SNc dopamine neurons to boost appetitive locomotion during predatory hunting. These results may provide some clues to the above unanswered questions. They supported the recent studies showing the involvement of SNc dopamine neurons in the vigor of body movements (Jin and Costa, 2010; Dodson et al., 2016; Howe and Dombeck, 2016; da Silva et al., 2018; Coddington and Dudman, 2018).

### More considerations on the functions of SC-SNc pathway

In their seminal studies, Redgrave and colleagues proposed that the SC-SNc pathway may serve as a route for salient visual stimuli to drive phasic activities of dopamine neurons (Dommett et al., 2005). In primate, this pathway may mediate visually-evoked reward expectation signals in dopamine neurons during reinforcement learning (Takakuwa et al., 2017). In the present study, we recorded single-unit activity of SNc-projecting SC neurons in head-fixed walking mice (Movie S1), and unexpectedly found that the SNc-projecting SC neurons encode locomotion speed (Figure 2). This observation prompted us to examine the role of the SC-SNc pathway in regulating locomotion during predatory hunting (Figure 3-7). Our data may have added another perspective for understanding the functions of the SC-SNc pathway. Although we did not systematically examine the sensory responses of the recorded neurons, we do not rule out the possibility that these neurons may respond to salient sensory stimuli (e.g. visual or vibrissal tactile stimuli). In future study, it will be interesting to explore whether the SC-SNc pathway can integrate both sensory and locomotion-related signals to dynamically modulate appetitive locomotion during hunting.

### The origin of locomotion-related signals of SNc-projecting SC neurons

Where do the locomotion-speed signals of the SNc-projecting SC neurons originate? Several motor-related brain areas (e.g. SNr, PPTg, and motor cortex) directly project to the SC and may provide motor signals to the SC (Comoli et al., 2012). This speculation was supported by a recent study showing that the projection from the SNr is the strongest among the above motor-related brain areas (Doykos et al., 2020). The axons of GABAergic SNr neurons terminate in the lateral part of deep layers of the SC (Kaneda et al., 2008), a region that contains SNc-projecting SC neurons studied here. The inhibition and excitation of SNr neurons well predict the initiation and suppression of locomotion, respectively (Freeze et al., 2013). These studies suggested that locomotion-related signals of SNc-projecting SC neurons may at least partially originate from the SNr, which is the primary output of basal ganglia.

## SUPPLEMENTARY INFORMATION

Supplementary information includes eight figures, seven movies and four tables.

## ACKNOWLEDGMENTS

We thank Drs. Thomas Südhof, Karl Deisseroth and Minmin Luo for providing plasmids and mouse lines. This work was supported by the National Natural Science Foundation of China (31925019 and 31671095 to P.C., 31771150 to Y.W., Top talent program of Hebei province to F.Z.), the open funds of the State Key Laboratory of Medical Neurobiology, and the Institutional Funding from NIBS. All data are archived in NIBS.

## AUTHOR CONTRIBUTIONS

P.C., C.S., J.Z., M.H., and F.Z. conceived the study. C.S., M.H., Q.P., and A.L. did injections and fiber implantation. C.S., Z.X., Q.P., H.G., and Y.X. did behavioral tests. C.S. did single-unit recording. M.H. did histological analyses. C.S. and Z.C. did slice physiology. Y.W., F.S., Y.L., J.Z., F.Z., M.H., and X.Q. provided reagents. D.L., C.S., Z.X., H.G., Z.C. and P.C. analyzed data. P.C. wrote the manuscript.

## MATERIALS AND METHODS

### Animals

All experimental procedures were conducted following protocols approved by the Administrative Panel on Laboratory Animal Care at the National Institute of Biological Sciences, Beijing (NIBS). The *vGlut2-IRES-Cre* (Vong et al., 2011), *GAD2-IRES-Cre* (Taniguchi et al., 2011), *DAT-IRES-Cre* (Backman et al., 2006), and *Ai14* (Madisen et al., 2010) mouse lines were imported from the Jackson Laboratory (JAX Mice and Services). Mice were maintained on a circadian 12-h light/12-h dark cycle with food and water available ad libitum. Mice were housed in groups (3–5 animals per cage) before they were separated 3 days prior to virus injection. After virus injection, each mouse was housed in one cage for 3 weeks before subsequent experiments. To avoid potential sex-specific differences, we used male mice only.

### AAV vectors

Two AAV serotypes (AAV-DJ, AAV2-retro) were used. The AAVs used in the present study are listed in Table S1. The viral particles were purchased from Shanghai Taitool Bioscience Inc. and Brain VTA Inc. The viral vector titers before dilution were in the range of 0.8-1.5×10^13^ viral particles/ml. The final titer used for AAV injection is 5×10^12^ viral particles/ml.

### Stereotaxic injection

Mice were anesthetized with an intraperitoneal injection of tribromoethanol (125–250 mg/kg). Standard surgery was performed to expose the brain surface above the superior colliculus (SC), substantia nigra pars compacta (SNc), ventral tegmental area (VTA), zona incerta (ZI) or dorsal striatum (DS). Coordinates used for SC injection were as follows: bregma −3.60 mm, lateral ± 1.30 mm, and dura −1.75 mm. Coordinates used for SNc injection were as follows: bregma −3.40 mm, lateral ± 1.25 mm, and dura −4.00 mm. Coordinates used for VTA injection were as follows: bregma −3.40 mm, lateral ± 0.50 mm, and dura −4.00 mm. Coordinates used for DS injection were as follows: bregma 0.74 mm, lateral ± 1.50 mm, and dura −2.40 mm. Coordinates used for ZI injection were: bregma −2.00 mm, lateral ± 1.25 mm and dura −4.25 mm. The AAVs and CTB were stereotaxically injected with a glass pipette connected to a Nano-liter Injector 201 (World Precision Instruments, Inc.) at a slow flow rate of 0.15 μl / min to avoid potential damage to local brain tissue. The pipette was withdrawn at least 20 min after viral injection.

For optogenetic activation and synaptic inactivation experiments, AAV injections were bilateral. For anterograde and retrograde tracing experiments, CTB injection was unilateral. Histological analyses were conducted one week (for CTB) and at least three weeks (for AAV) after injection. Experimental designs related to viral injection are summarized in Table S2.

### Optical fiber implantation

Thirty minutes after AAV injection, a ceramic ferrule with an optical fiber (230 µm in diameter, numerical aperture = 0.37) was implanted with the fiber tip on top of the SNc (bregma −3.40 mm, lateral ± 1.25 mm, dura −3.80 mm) or the dorsal striatum (bregma 0.74 mm, lateral ± 1.50 mm, and dura −2.20 mm). The ferrule was then secured on the skull with dental cement. After implantation, the skin was sutured, and antibiotics were applied to the surgical wound. The optogenetic experiments were conducted 3 weeks after optical fiber implantation. All experimental designs related to optical fiber implantation are summarized in Table S2. For optogenetic stimulation, the output of the laser was measured and adjusted to 2, 5, 10, 15 and 20 mW before each experiment. The pulse onset, duration, and frequency of light stimulation were controlled by a programmable pulse generator attached to the laser system.

### Single-unit recording

Antidromic activation strategy was used to identify the single-unit activity of SNc-projecting SC neurons. AAV-hSyn-ChR2-mCherry was injected into the SC of wild-type mice, followed by an optical fiber implanted above the SNc. Three weeks after viral injection, single-unit recording was performed with a tungsten electrode in the SC of head-fixed awake mouse. The tungsten electrode was vertically advanced into the lateral SC with a Narishige micro-manipulator. The spikes were amplified by a differential amplifier (Model 1800, A-M Systems, Everett, WA, USA), digitized (10 kHz) and stored by Spike2 software (Version 7.03). When the single-unit activity was isolated, we tested if the units were from SNc-projecting SC neurons. The putative SNc-projecting SC neurons were identified by the antidromic spikes evoked by light-pulses (473 nm, 1 ms, 2 mW) illuminating ChR2-mCherry+ axon terminals in the SNc. The antidromically evoked spikes had to conform to two criteria: first, their latency to the light pulse should be less than 5 ms; second, their waveform should be similar to that of spikes evoked by locomotion (Figure 2B). Only units with spikes faithfully following the light stimulations with latency less than 5 ms were further tested for locomotion-correlated activity (Figure 2C). The spike sorting was performed with Spike2 Software (Version 7.03). For a certain train of action potential, after setting the threshold of the spikes, Spike2 automatically generated the templates and performed the spike-sorting. The quality of spike clustering was further confirmed by principal component analysis (Figure S4B). The single-unit activity of SNc-projecting SC units was recorded with simultaneously measuring the instantaneous locomotion speed of mice walking on the treadmill (Nanjing Thinktech Inc.).

### Verification of recording sites

The recording sites of the putative SNc-projecting SC neurons were marked with electrolytic lesions applied by passing positive currents (40 µA, 10 s) through the tungsten electrode. Under deep anesthesia with urethane, the brain was perfused with saline and PBS containing 4% PFA. After regular histological procedure, frozen sections were cut at 40 µm in thickness and counterstained with DAPI for histological verification of recording sites.

### Preparation of behavioral tests

After AAV injection and fiber implantation, the mice were housed individually for 3 weeks before the behavioral tests. Before the behavioral tests, they were handled daily by the experimenters for at least 3 days. On the day of the behavioral test, the mice were transferred to the testing room and were habituated to the room conditions for 3 h before the experiments started. The apparatus was cleaned with 20% ethanol to eliminate odor cues from other mice. All behavioral tests were conducted during the same circadian period (13:00–19:00). All behaviors were scored by the experimenters, who were blind to the animal treatments.

### Behavioral paradigm for predatory hunting

The procedure of predatory hunting experiment was described previously (Shang et al., 2019). Before the predatory hunting test, the mice went through a 9-day habituation procedure (Day H1–H9). On each of the first three habituation days (Day H1, H2, H3), three cockroaches were placed in the home-cage (with standard chow) of mice at 2:00 PM. The mice readily consumed the cockroaches within 3 h after cockroach appearance. On Day H3, H5, H7, and H9, we initiated 24-h food deprivation at 7:00 PM by removing chow from the home-cage. On Day H4, H6, and H8 at 5:00 PM, we let the mice freely explore the arena (25 cm x 25 cm) for 10 min, followed by three trials of hunting practice for the cockroach. After hunting practice, we put the mice back in their home-cages and returned the chow at 7:00 PM. On the test day, we let the mice freely explore the arena for 10 min, followed by three trials of predatory hunting. After the tests, the mice were put back in their home-cage, followed by the return of chow. The cockroach was purchased from a merchant in Tao-Bao Online Stores (www.taobao.com).

Before the hunting practice or test, the mice were transferred to the testing room and habituated to the room conditions for 3 h before the experiments started. The arena was cleaned with 20% ethanol to eliminate odor cues from other mice. All behaviors were scored by the experimenters, who were blind to the animal treatments. Hunting behaviors were measured in an arena (20 cm × 20 cm, square open field) without regular mouse bedding. After entering, the mice explored the arena for 10 min, followed by the introduction of a cockroach. For each mouse, predatory hunting was repeated for three trials. Each trial began with the introduction of prey to the arena. The trial ended when the predator finished ingesting the captured prey. After the mice finished ingesting the prey body, debris was removed before the new trial began.

### Measurement of appetitive locomotion and predatory attack in predatory hunting

In the paradigm of predatory hunting, mouse behavior was recorded in the arena with three orthogonally positioned cameras (50 frames/sec; Point Grey Research, Canada). With the video taken by the overhead camera, the instantaneous head orientation of predator relative to prey (azimuth angle) and predator-prey distance (PPD) was analyzed with the Software EthoVision XT 14 (Noldus Information Technology). The episode of approach was identified by two empirical criteria (Hoy et al., 2016). First, the PPD should continuously decrease until it is less than 3 cm. Second, the azimuth angle of mouse head to cockroach should be within the range of −90 deg to +90 deg. In WT mice, each trial of predatory hunting contains 10∼20 episodes of approach. Speed of approach and frequency of approach were used to quantitatively measure the appetitive locomotion in the episodes of approach. Speed of approach of each mouse was calculated by averaging the peak speed in all the approach episodes in the trial. Frequency of approach was the total number of approach episodes divided by the time to prey capture in the trial.

With the videos taken by the two horizontal cameras, we carefully identified predatory attacks with jaw by replaying the video frame by frame (50 frames/sec). We marked the predatory jaw attacks with yellow vertical lines in the behavioral ethogram of predatory hunting. With this method, we measured three parameters of predatory hunting: time to prey capture, latency to jaw attack, and frequency of jaw attack. Time to prey capture was defined as the time between the introduction of prey and the last jaw attack. Latency to jaw attack was defined as the time between the introduction of the prey and the first jaw attack from the predator. Frequency of jaw attack was defined as the number of jaw attacks divided by time to prey capture. Data for three trials were averaged.

### Measurement of defensive locomotion triggered by looming visual stimuli

Measurement of defensive locomotion triggered by looming visual stimulus was described previously (Shang et al., 2018). Briefly, defensive locomotion was measured in an arena (35 cm × 35 cm, square open field) with corn-cob bedding. No shelter was provided. A regular computer monitor was positioned above the arena for presentation of overhead looming visual stimuli. After entering, the mice explored the arena for 10 min. This was followed by the presentation of three cycles of overhead looming visual stimuli consisting of an expanding dark disk. The visual angel of the dark disk was expanded from 2 to 20 degrees within 250 ms. Luminance of the dark disk and background were 0.1 and 3.6 cd/m^2^, respectively. Mouse behavior was recorded (50 frames/sec; Point Grey Research, Canada) by two orthogonally positioned cameras with LEDs providing infrared illumination. The instantaneous location of the mouse in the arena was measured by a custom-written Matlab program. The instantaneous locomotion speed was calculated with a 200 ms time-bin. The Matlab code is available upon request.

### Measurement of locomotion in linear runway

Mouse behavior was recorded in the linear runway (10 cm x 16 cm x 120 cm) with an overhead camera (50 frames/sec; Point Grey Research, Canada). With the video taken by the overhead camera, we measured the instantaneous locomotion speed with the Software EthoVision XT 14 (Noldus Information Technology).

### Slice physiological recording

Preparation of acute brain slices was performed according to the published work (Liu et al., 2017). Brain slices containing the SC or SNc were prepared from adult mice anesthetized with isoflurane before decapitation. Brains were rapidly removed and placed in ice-cold oxygenated (95% O_2_ and 5% CO_2_) cutting solution (228 mM sucrose, 11 mM glucose, 26 mM NaHCO_3_, 1 mM NaH_2_PO_4_, 2.5 mM KCl, 7 mM MgSO_4_, and 0.5 mM CaCl_2_). Coronal brain slices (400 μm) were cut using a vibratome (VT 1200S, Leica Microsystems, Wetzlar, Germany). The slices were incubated at 28°C in oxygenated artificial cerebrospinal fluid (ACSF: 125 mM NaCl, 2.5 mM KCl, 1.25 mM NaH_2_PO_4_, 1.0 mM MgCl_2_, 25 mM NaHCO_3_, 15 mM glucose, and 2.0 mM CaCl_2_) for 30 min (∼305 mOsm, pH 7.4). The slices were then kept at room temperature under the same conditions for 30 min before transfer to the recording chamber at room temperature. The ACSF was perfused at 1 ml/min. The acute brain slices were visualized with a 40× Olympus water immersion lens, differential interference contrast (DIC) optics (Olympus Inc., Japan), and a CCD camera.

Patch pipettes were pulled from borosilicate glass capillary tubes (Cat #64-0793, Warner Instruments, Hamden, CT, USA) using a PC-10 pipette puller (Narishige Inc., Tokyo, Japan). For recording of postsynaptic currents (voltage clamp), pipettes were filled with solution (126 mM Cs-methanesulfonate, 10 mM HEPES, 1 mM EGTA, 2 mM QX-314 chloride, 0.1 mM CaCl_2_, 4 mM Mg-ATP, 0.3 mM Na-GTP, 8 mM Na-Phosphocreatine, pH 7.3 adjusted with CsOH, ∼290 mOsm) (Kim et al., 2015). For recording of action potentials (current clamp), pipettes were filled with solution (135 mM K-methanesulfonate, 10 mM HEPES, 1 mM EGTA, 1 mM Na-GTP, 4 mM Mg-ATP, pH 7.4). The resistance of pipettes varied between 3.0–3.5 MΩ. The current and voltage signals were recorded with MultiClamp 700B and Clampex 10 data acquisition software (Molecular Devices). After establishment of the whole-cell configuration and equilibration of the intracellular pipette solution with the cytoplasm, series resistance was compensated to 10–15 MΩ. Recordings with series resistances of > 15 MΩ were rejected. An optical fiber (230 μm in diameter) was used to deliver light pulses, with the fiber tip positioned 500 μm above the brain slices. Laser power was adjusted to 2, 5, 10, or 20 mW. Light-evoked action potentials from ChR2-mCherry+ neurons in the SC were triggered by a light-pulse train (473 nm, 2 ms, 10 Hz, 20 mW) synchronized with Clampex 10 data acquisition software (Molecular Devices). Light-evoked postsynaptic currents from SNc neurons were triggered by single light pulses (2 ms) in the presence of 4-aminopyridine (4-AP, 20 μM) and tetrodotoxin (TTX, 1 μM). D-AP5 (50 μM)/CNQX (20 μM) or picrotoxin (PTX, 50 μM) were perfused with ACSF to examine the neurotransmitter/receptor type of optically-evoked postsynaptic currents.

### Fiber photometry

A fiber photometry system (ThinkerTech, Nanjing, China) was used for recording GRAB_DA_ signals from genetically identified neurons (Sun et al., 2018). To induce fluorescence signals, a laser beam from a laser tube (488 nm) was reflected by a dichroic mirror, focused by a 10× lens (N.A. 0.3) and coupled to an optical commutator. A 2-m optical fiber (230 μm in diameter, N.A. 0.37) guided the light between the commutator and implanted optical fiber. To minimize photo bleaching, the power intensity at the fiber tip was adjusted to 0.02 mW. The GRAB_DA_ fluorescence was band-pass filtered (MF525-39, Thorlabs) and collected by a photomultiplier tube (R3896, Hamamatsu). An amplifier (C7319, Hamamatsu) was used to convert the photomultiplier tube current output to voltage signals, which were further filtered through a low-pass filter (40 Hz cut-off; Brownlee 440). The analogue voltage signals were digitalized at 100 Hz and recorded by a Power 1401 digitizer and Spike2 software (CED, Cambridge, UK).

AAV-DJ-hSyn-GRAB-DA was stereotaxically injected into the dorsal striatum of *WT* mice followed by optical fiber implantation above the injected site (see “Stereotaxic injection” and “Optical fiber implantation”). Two weeks after AAV injection, fiber photometry was used to record GRAB-DA signals from the cell bodies of dorsal striatum neurons in freely moving mice. A flashing LED triggered by a 1-s square-wave pulse was simultaneously recorded to synchronize the video and GRAB-DA signals. For recordings from freely moving mice, mice with optical fibers connected to the fiber photometry system freely explored the arena for 10 min. After the experiments, the optical fiber tip sites above the dorsal striatum neurons were histologically examined in each mouse.

### Histological procedures

Mice were anesthetized with isoflurane and sequentially perfused with saline and phosphate buffered saline (PBS) containing 4% paraformaldehyde (PFA). Brains were removed and incubated in PBS containing 30% sucrose until they sank to the bottom. Post-fixation of the brain was avoided to optimize immunohistochemistry of GABA and glutamate. Cryostat sections (40 μm) containing the SC, SNc or DS were collected, incubated overnight with blocking solution (PBS containing 10% goat serum and 0.7% Triton X-100), and then treated with primary antibodies diluted with blocking solution for 3–4 h at room temperature. Primary antibodies used for immunohistochemistry are displayed in Table S1. Primary antibodies were washed three times with washing buffer (PBS containing 0.7% Triton X-100) before incubation with secondary antibodies (tagged with Cy2, Cy3, or Cy5; dilution 1:500; Life Technologies Inc., USA) for 1 h at room temperature. Sections were then washed three times with washing buffer, stained with DAPI, and washed with PBS, transferred onto Super Frost slides, and mounted under glass coverslips with mounting media.

Sections were imaged with an Olympus (Japan) VS120 epifluorescence microscope (10× objective lens) or an Olympus FV1200 laser scanning confocal microscope (20× and 60× oil-immersion objective lens). Samples were excited by 488, 543, or 633 nm lasers in sequential acquisition mode to avoid signal leakage. Saturation was avoided by monitoring pixel intensity with Hi-Lo mode. Confocal images were analyzed with ImageJ software.

### Quantification of synaptic puncta density

The micrographs used for measuring puncta density of SynaptoTag (Figure 1, B and D) were acquired with a 63× objective of Zeiss confocal system and analyzed with NIH Image J. The analysis of the synaptic puncta was described previously (Cao et al., 2013). In brief, the scale of micrographs was set in NIH Image J based on the physical dimension of micrographs acquired by Zeiss confocal system. After converting the micrographs from RGB color mode to 16-bit mode, the puncta in micrographs were binarized and the puncta density was measured automatically by NIH Image J. Then the puncta density in the SNc of each mouse was normalized by dividing with that in the intermediate layer of the lateral SC (Figure 1E).

### Cell-counting strategies

Cell-counting strategies are summarized in Table S3. For counting cells in the SC, we collected 40-μm coronal sections from bregma −3.28 to bregma −4.48 for each mouse. Six sections evenly spaced by 200 μm were sampled for immunohistochemistry to label cells positive for different markers. We acquired micrographs (10× objective, Olympus FV1200 microscope, Japan) within intermediate and deep layers of the SC followed by cell counting with ImageJ software. We calculated the percentages of glutamate+ and GABA+ neurons in the neuronal population retrogradely labeled by CTB-555. For counting cells in the SNc, we collected coronal sections (40 μm) from bregma −2.80 to bregma −3.80 for each mouse. Five sections evenly spaced by 200 μm were sampled for immunohistochemistry to label SNc cells positive for different markers. After image acquisition, we outlined the SC and SNc followed by cell counting with ImageJ software. The boundary of SNc in coronal sections was identified based on TH staining.

## DATA QUANTIFICATION AND STATISTICAL ANALYSIS

All experiments were performed with anonymized samples in which the experimenter was unaware of the experimental conditions of the mice. For the statistical analyses of experimental data, Student t-test and One-Way ANOVA were used. The “n” used for these analyses represents number of mice or cells. See the detailed information of statistical analyses in figure legend and in Table S4.

## DATA AND CODE AVAILABILITY

The data that support the findings of this study are available from the corresponding author upon reasonable request. The MATLAB code for data analyses is available from the corresponding author upon request.

## Supplementary Materials

**Figure S1.**
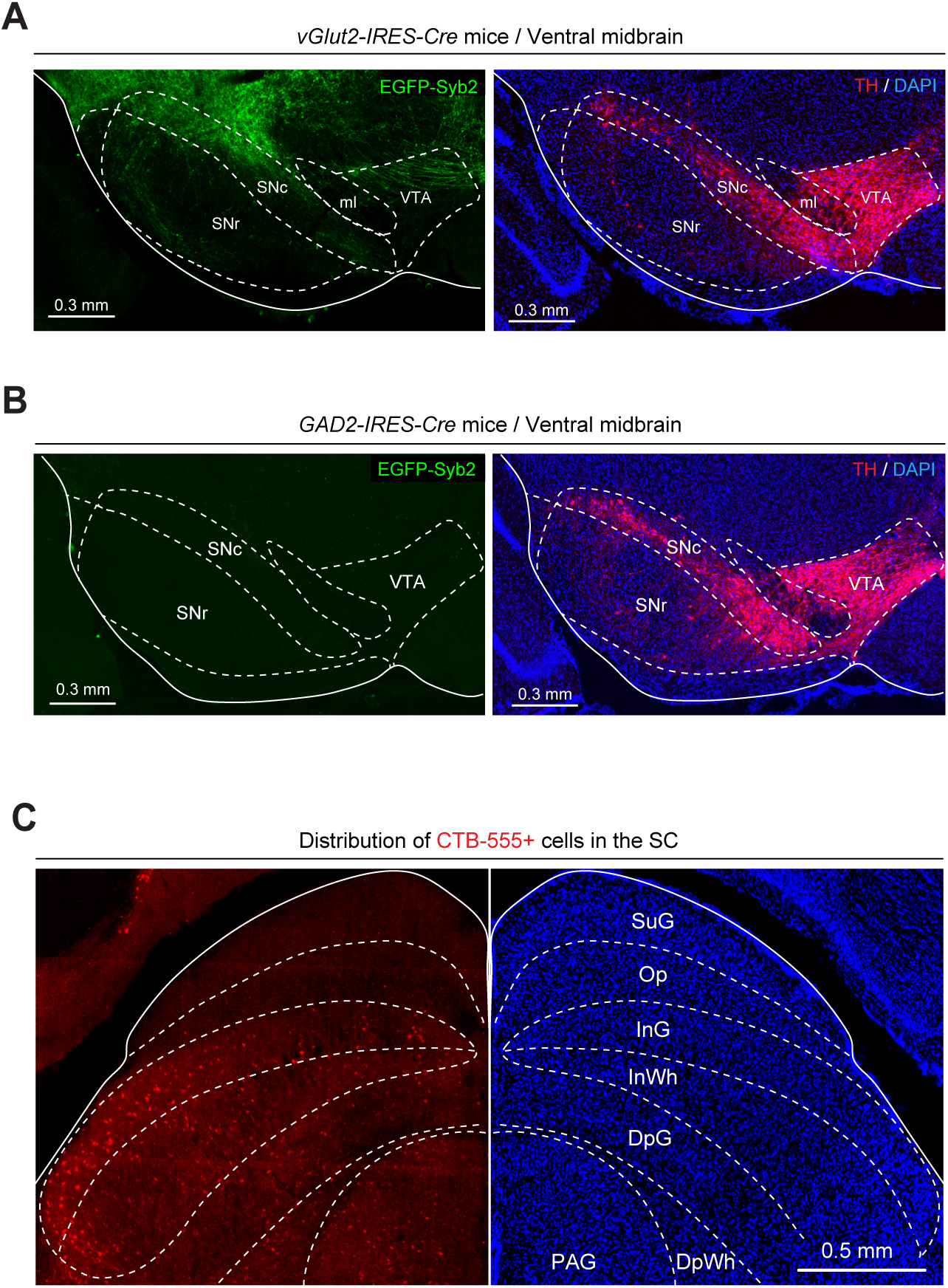
Cell-type-specific mapping of tectonigral pathway. (Related to Figure 1) **(A, B)** Single-channel and merged micrographs showing EGFP-Syb2+ axon terminals in the ventral midbrain of *vGlut2-IRES-Cre* (A) and *GAD2-IRES-Cre* mice (B). The boundaries of SNc and VTA were determined by immunostaining of tyrosine hydroxylase (TH, red) for dopamine neurons. (**C**) An example coronal section of the SC showing the distribution of SNc-projecting SC neurons that were labeled by CTB-555. Scale bars were indicated in the graphs.

**Figure S2.**
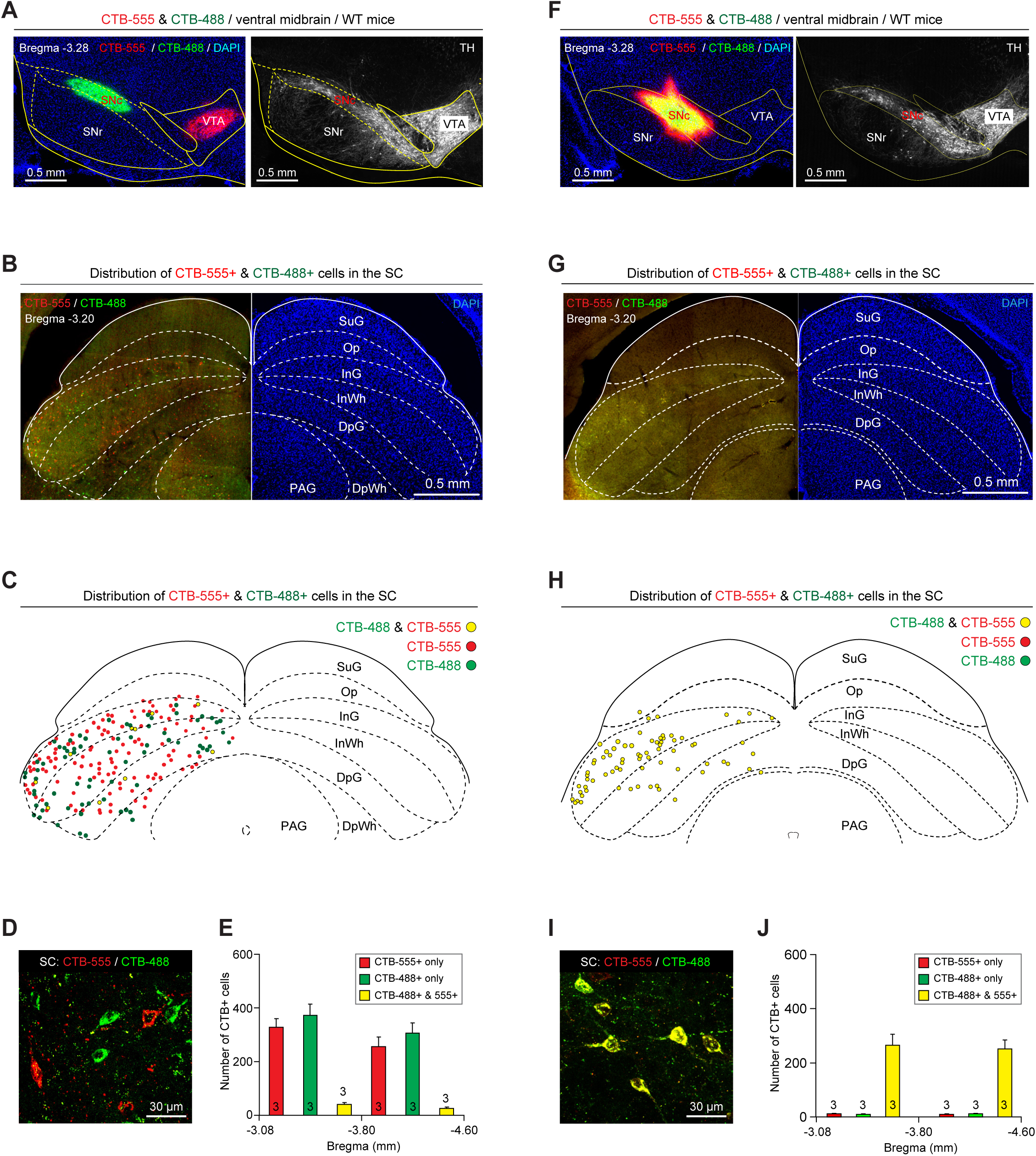
SC-SNc pathway and SC-VTA pathway are anatomically segregated. (Related to Figure 1) **(A)** Example coronal section of the ventral midbrain showing injection of CTB-488 and CTB-555 into the SNc and VTA (*left*), the boundaries of which were delineated according to the immunofluorescence of TH (*right*). **(B, C)** An example coronal section of the SC (B) and the corresponding illustration (C) showing the distribution of CTB-555+ & CTB-488+ cells in the SC. **(D, E)** Example micrograph (D) and quantitative analyses (E) showing CTB-555+ & CTB-488+ SC neurons were largely segregated. (**F**) Example coronal section of the ventral midbrain showing injection of mixed CTB-488 & CTB-555 into the SNc (*left*), the boundary of which was delineated according to the immunofluorescence of TH (*right*). (**G, H**) An example coronal section of the SC (G) and the corresponding illustration (H) showing the distribution of CTB-555+ & CTB-488+ cells in the SC. (**I, J**) Example micrograph (I) and quantitative analyses (J) showing the retrogradely labeled SC neurons are mostly positive for both CTB-555 and CTB-488. Numbers of mice (E and J) are indicated in the graphs. Data in (E and J) are means ± SEM. Scale bars are labeled in the graphs.

**Figure S3.**
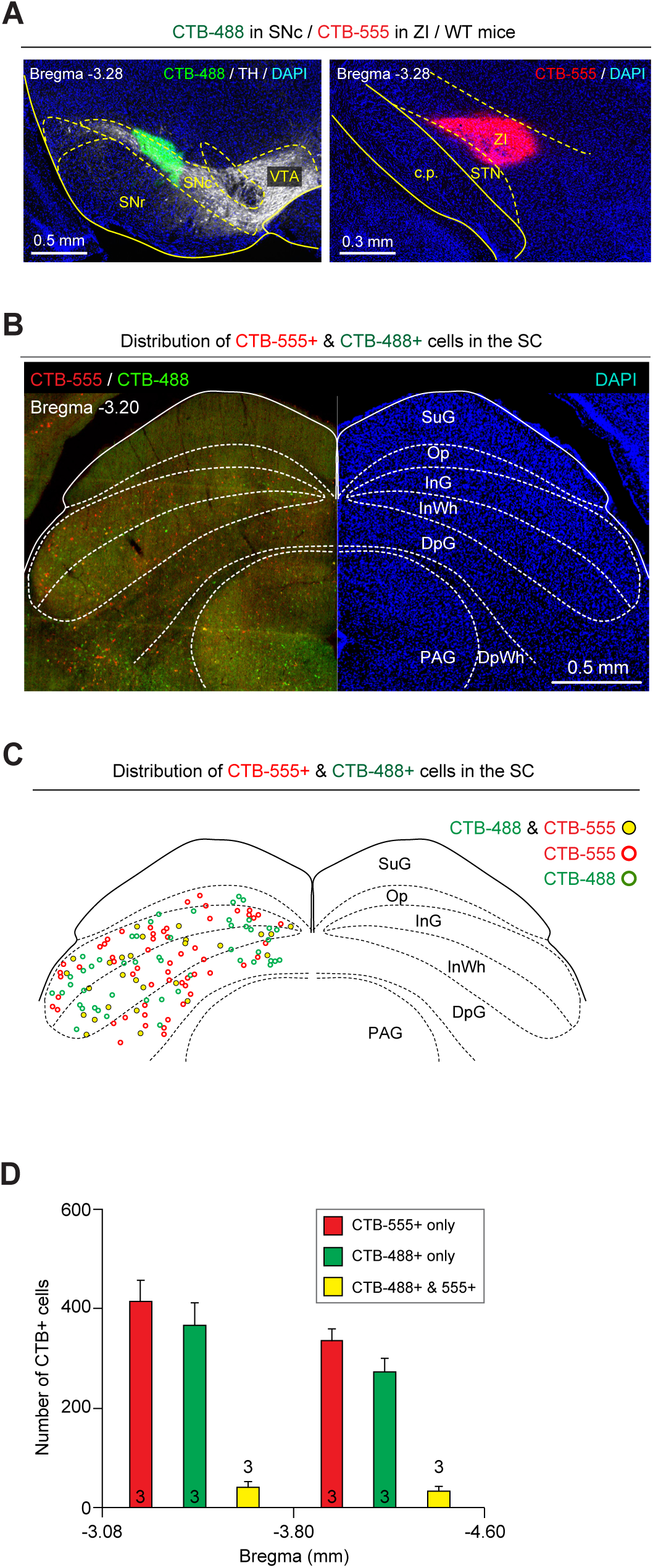
SC-SNc pathway and SC-ZI pathway are anatomically segregated. (Related to Figure 1) **(A)** Example coronal brain section showing injection of CTB-488 and CTB-555 into the SNc (*left*) and ZI (*right*), respectively. The boundary of SNc was delineated according to the immunofluorescence of TH (*left*). **(B, C)** An example coronal brain section (B) and the corresponding illustration (C) showing the distribution of CTB-555+ & CTB-488+ cells in the SC. **(D)** Quantitative analyses showing CTB-555+ & CTB-488+ cells in the SC were largely segregated. Numbers of mice (D) are indicated in the graphs. Data in (D) are means ± SEM. Scale bars are labeled in the graphs.

**Figure S4.**
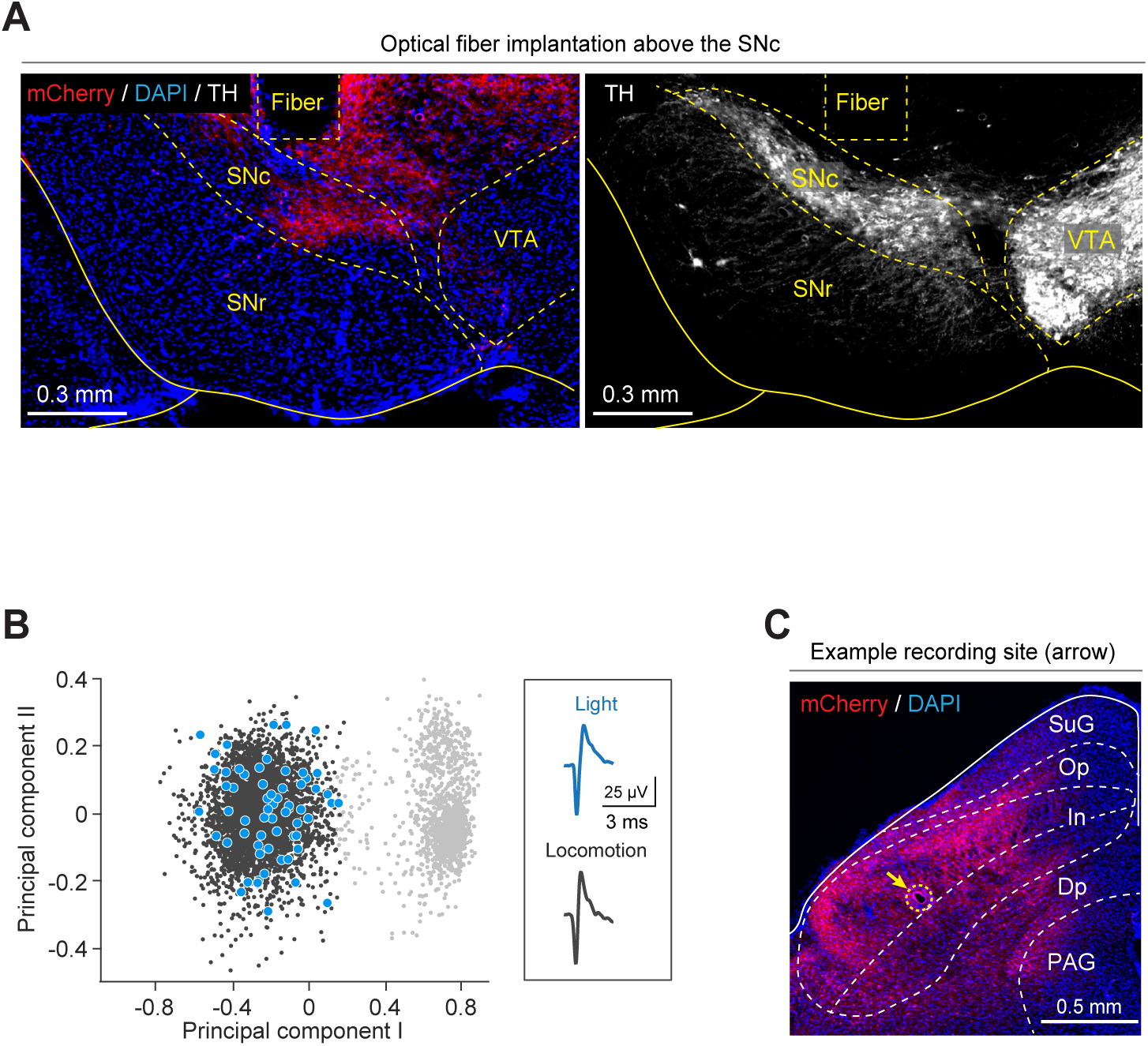
SNc-projecting SC neurons encode the speed of locomotion. (Related to Figure 2) (**A**) Example coronal section of the ventral midbrain showing the optical-fiber track above ChR2-mCherry^+^ axon terminals in the SNc (*left*), the boundary of which was determined by immunofluorescence of TH for dopamine neurons (*right*). **(B)** Principal component analyses of light-evoked spikes (blue) and locomotion-evoked spikes (black) of an example putative SNc-projecting SC neuron. Gray dots, noise. **(C)** Example coronal section of the SC showing a recording site marked by electrolytic lesion (arrow) in the intermediate layer (In) of the SC. Scale bars are labeled in the graphs.

**Figure S5.**
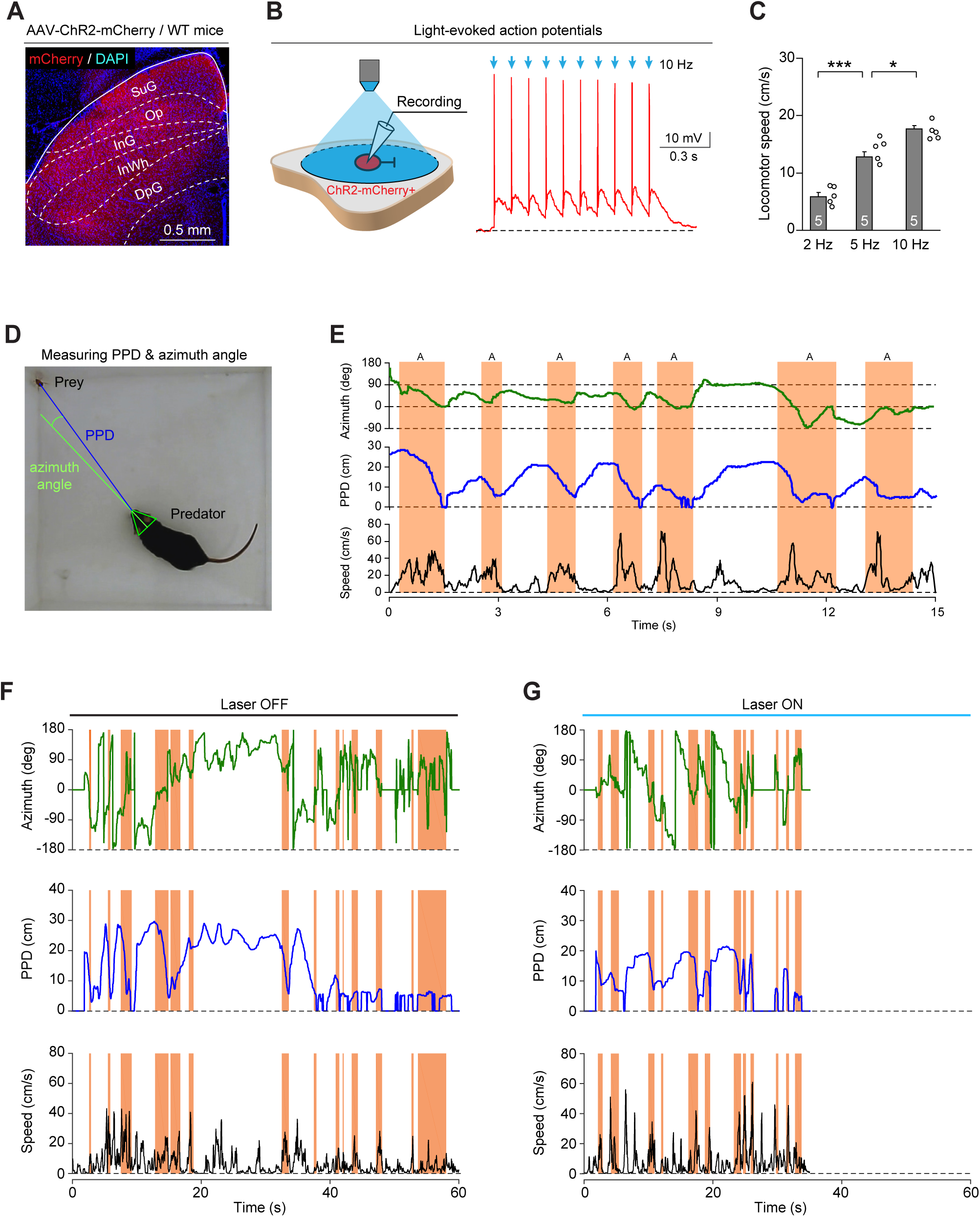
Activation of the SC-SNc pathway promotes appetitive locomotion during predatory hunting. (Related to Figure 3) **(A)** Example coronal section of the SC showing ChR2-mCherry expression in the SC of WT mice. (**B**) Schematic diagram (*left*) and example trace (*right*) showing light pulses (2 ms, 473 nm, 10 Hz, 10 pulses) reliably triggered action potential firing from ChR2-mCherry^+^ SC neurons in acute SC slices. **(C)** Quantitative analyses of average locomotion speed of mice in the linear runway during photostimulation of the SC-SNc pathway (10 ms, 6 s, 10 mW) with different frequencies (2 Hz, 5 Hz, 10 Hz). **(D)** An example picture showing computer-aided measurement of azimuth angle and prey-predator distance (PPD) when predator approached prey. **(E)** Aligned time courses of azimuth angle (*top*), prey-predator distance (PPD, *middle*), and locomotion speed (*bottom*) of an example mouse in predatory hunting, showing the identification of approach episodes (shaded areas in orange). The intermittent approach episodes were characterized by azimuth within a narrow range (−90–90 deg), by decreased PPD and by pulses of locomotion speed. For the detailed criteria to identify the approach episodes, see Methods. (**F, G**) Time courses of azimuth angle (*top*), PPD (*middle*), and locomotion speed (*bottom*) during predatory hunting of an example mouse without (F, Laser OFF) and with (G, Laser ON) photostimulation of the SC-SNc pathway. Numbers of mice (C) are indicated in the graphs. Data in (C) are means ± SEM. Statistic analyses (C) were performed using Student t-test (*** P<0.001; * P<0.05). For the P values, see Table S4. Scale bars are labeled in the graphs.

**Figure S6.**
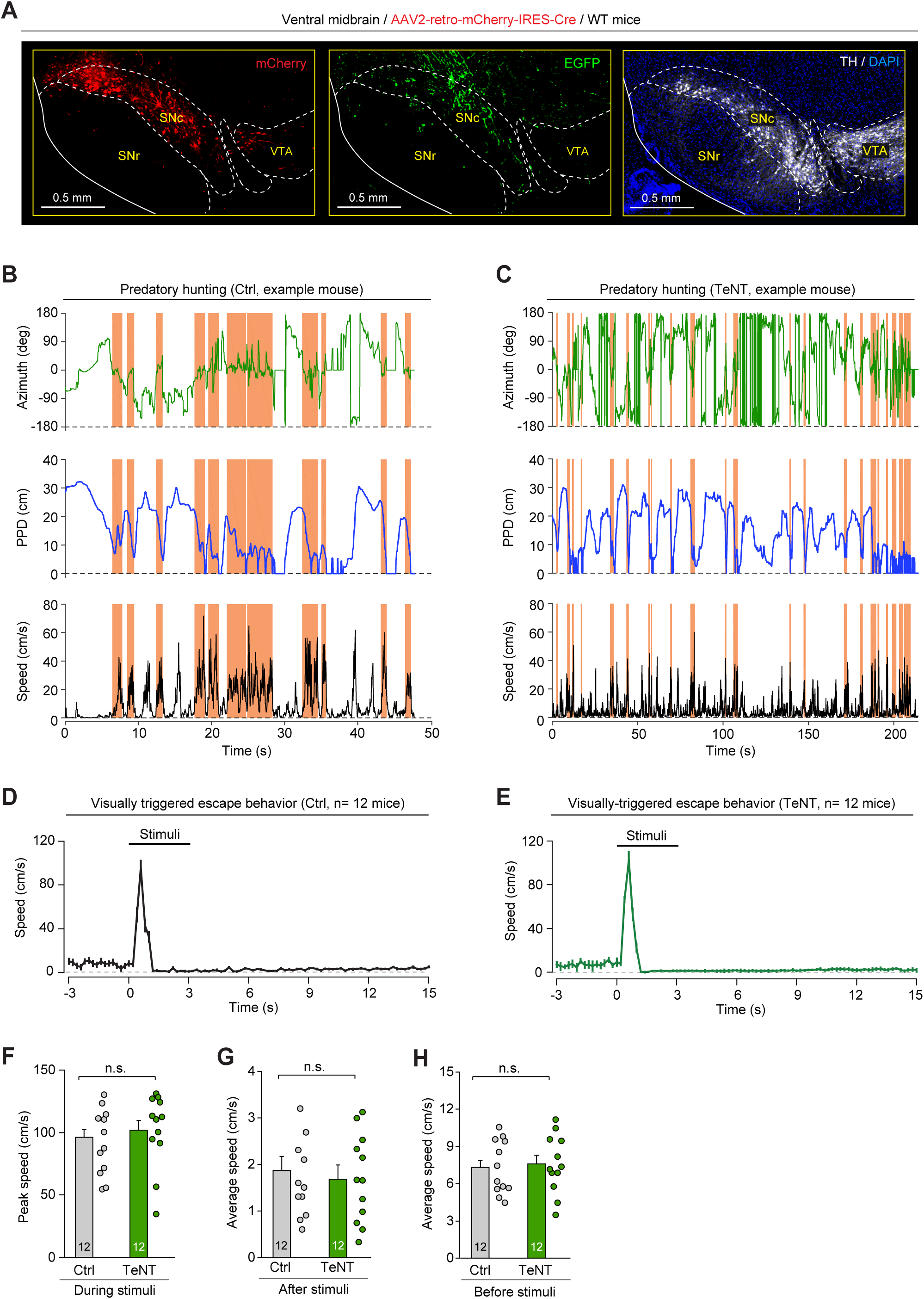
The SNc-projecting SC neurons are required for appetitive locomotion during predatory hunting. (Related to Figure 4) **(A)** Example coronal section of ventral midbrain showing mCherry+ cells in the SNc (*left*) are intermingled with EGFP+ axon terminals from SC neurons (*middle*). The boundary of the SNc is determined by immunofluorescence of TH (*right*). (**B, C**) Time courses of azimuth angle (*top*), PPD (*middle*), and locomotion speed (*bottom*) during predatory hunting of example mice without (B, Ctrl) and with (C, TeNT) synaptic inactivation of the SNc-projecting SC neurons. **(D, E)** Time courses of locomotion speed before, during and after looming visual stimuli in mice without (Ctrl, D) and with (TeNT, E) synaptic inactivation of SNc-projecting SC neurons. (**F-H**) Quantitative analyses of peak locomotion speed during stimuli (F), average locomotion speed after stimuli (G), and average locomotion speed before stimuli (H) of mice without (Ctrl) and with (TeNT) synaptic inactivation of SNc-projecting SC neurons. Scale bars are labeled in the graphs (A). Numbers of mice are indicated in the graphs (D-H). Data in (D-H) are means ± SEM. Statistic analyses (F-H) are performed using Student t-test (n.s. P>0.1). For the P values, see Table S4.

**Figure S7.**
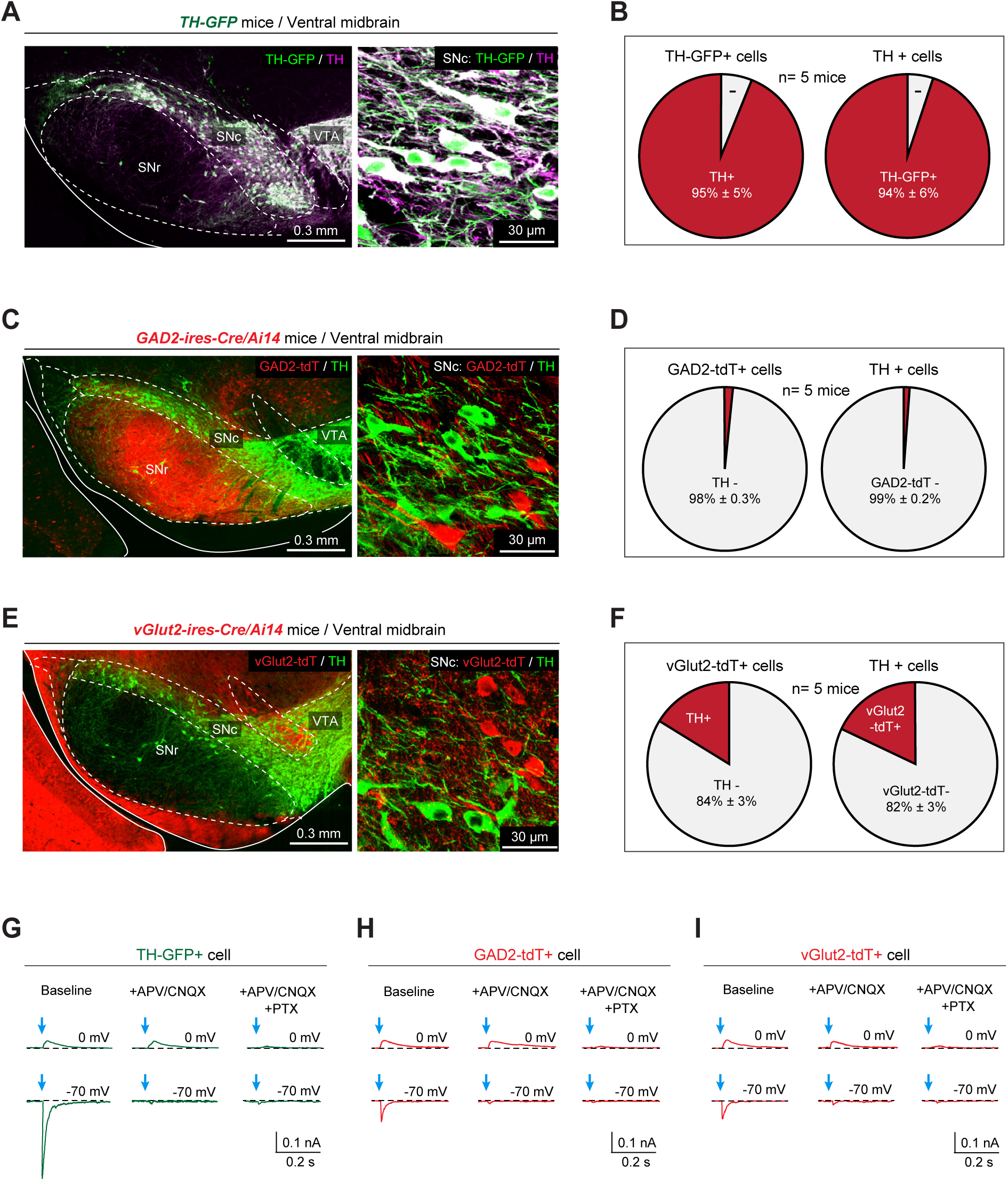
Dopamine neurons are the primary postsynaptic target of the SC-SNc pathway. (Related to Figure 5) **(A, B)** Example micrographs (A) and quantitative analyses (B) showing that TH-GFP+ cells and TH+ cells are largely overlapped in the SNc of *TH-GFP* mice. **(C, D)** Example micrographs (C) and quantitative analyses (D) showing that GAD2-tdT+ cells and TH+ cells are largely segregated in the SNc of *GAD2-IRES-Cre/Ai14* mice. **(E, F)** Example micrographs (E) and quantitative analyses (F) showing that vGlut2-tdT+ cells and TH+ cells are largely segregated in the SNc of *vGlut2-IRES-Cre/Ai14* mice. (**G-I**) Example traces of oIPSCs (*top*) and oEPSCs (*bottom*) from putative SNc dopamine neurons (TH-GFP+) (G), putative GAD2+ neurons (GAD2-tdT+) (H) and putative vGlut2+ neurons (vGlut2-tdT+) (I) with and without perfusion of antagonists of glutamate receptors (APV/CNQX) and GABAa receptor (PTX). Numbers of mice (B, D, F) are indicated in the graphs. Data in (B, D, F) are means ± SEM. Scale bars are labeled in the graphs.

**Figure S8.**
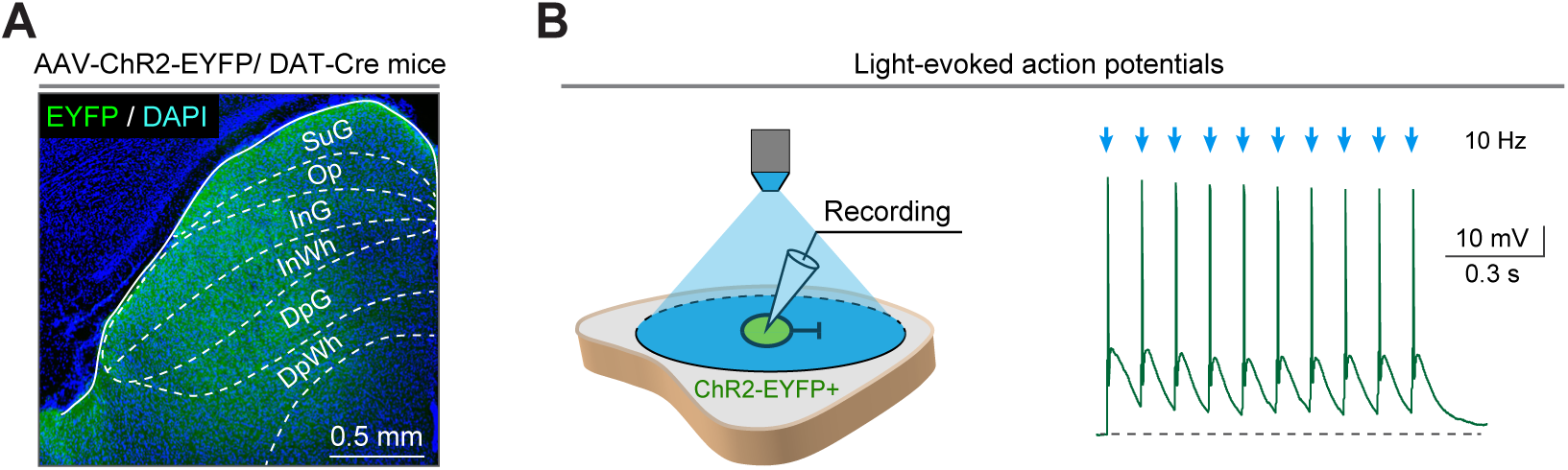
SNc dopamine neurons mediate the effect of SC-SNc pathway activation. (Related to Figure 7) (**A**) Example coronal section showing ChR2-EYFP expression in the SC. (**B**) Schematic diagram (*left*) and example trace (*right*) showing light pulses (2 ms, 473 nm, 10 Hz, 10 pulses) reliably triggered action potential firing from ChR2-EYFP+ SC neurons in acute SC slices. Scale bars are labeled in the graphs.

---

**Movie S1 Action potential firing of SNc-projecting SC neurons during locomotion**

This movie shows that the action potential firing of a putative SNc-projecting SC neuron recorded with an optrode is correlated with locomotion. The action potentials have been sorted and their waveforms are displayed in the corner of the movie.

**Movie S2 Predatory hunting of an example control mouse without photostimulation of the SC-SNc pathway**

This movie shows behavioral analyses of predatory hunting of an example control mouse without photostimulation of the SC-SNc pathway. The left part of the screen displays the video taken by the overhead camera in parallel with computer-aided analyses of azimuth angle and PPD in real-time. The right part of the screen displays the time courses of azimuth angle, locomotion speed and PPD during predatory hunting in real-time. The approach episodes were labeled with shaded areas in orange.

**Movie S3 Predatory hunting of an example test mouse with photostimulation of the SC-SNc pathway**

This movie shows behavioral analyses of predatory hunting of an example test mouse with photostimulation of the SC-SNc pathway. The left part of the screen displays the video taken by the overhead camera in parallel with computer-aided analyses of azimuth angle and PPD in real-time. The right part of the screen displays the time courses of azimuth angle, PPD and locomotion speed during predatory hunting in real-time. The approach episodes were labeled with shaded areas in orange.

**Movie S4 Predatory hunting of an example control mouse without synaptic inactivation of SNc-projecting SC neurons**

This movie shows behavioral analyses of predatory hunting of an example control mouse without synaptic inactivation of SNc-projecting SC neurons. The left part of the screen displays the video taken by the overhead camera in parallel with computer-aided analyses of azimuth angle and PPD in real-time. The right part of the screen displays the time courses of azimuth angle, locomotion speed and PPD during predatory hunting in real-time. The approach episodes were labeled with shaded areas in orange.

**Movie S5 Predatory hunting of an example test mouse with synaptic inactivation of SNc-projecting SC neurons**

This movie shows behavioral analyses of predatory hunting of an example test mouse with synaptic inactivation of SNc-projecting SC neurons. The left part of the screen displays the video taken by the overhead camera in parallel with computer-aided analyses of azimuth angle and PPD in real-time. The right part of the screen displays the time courses of azimuth angle, locomotion speed and PPD during predatory hunting in real-time. The approach episodes were labeled with shaded areas in orange.

**Movie S6 Defensive locomotion of an example control mouse without synaptic inactivation of SNc-projecting SC neurons**

This movie shows the overhead looming visual stimuli triggered escape followed by long-lasting freezing in an example control mouse without synaptic inactivation of SNc-projecting SC neurons.

**Movie S7 Defensive locomotion of an example test mouse with synaptic inactivation of SNc-projecting SC neurons**

This movie shows the overhead looming visual stimuli evoked immediate escape followed by long-lasting freezing in an example test mouse with synaptic inactivation of SNc-projecting SC neurons.

**Table S1.**
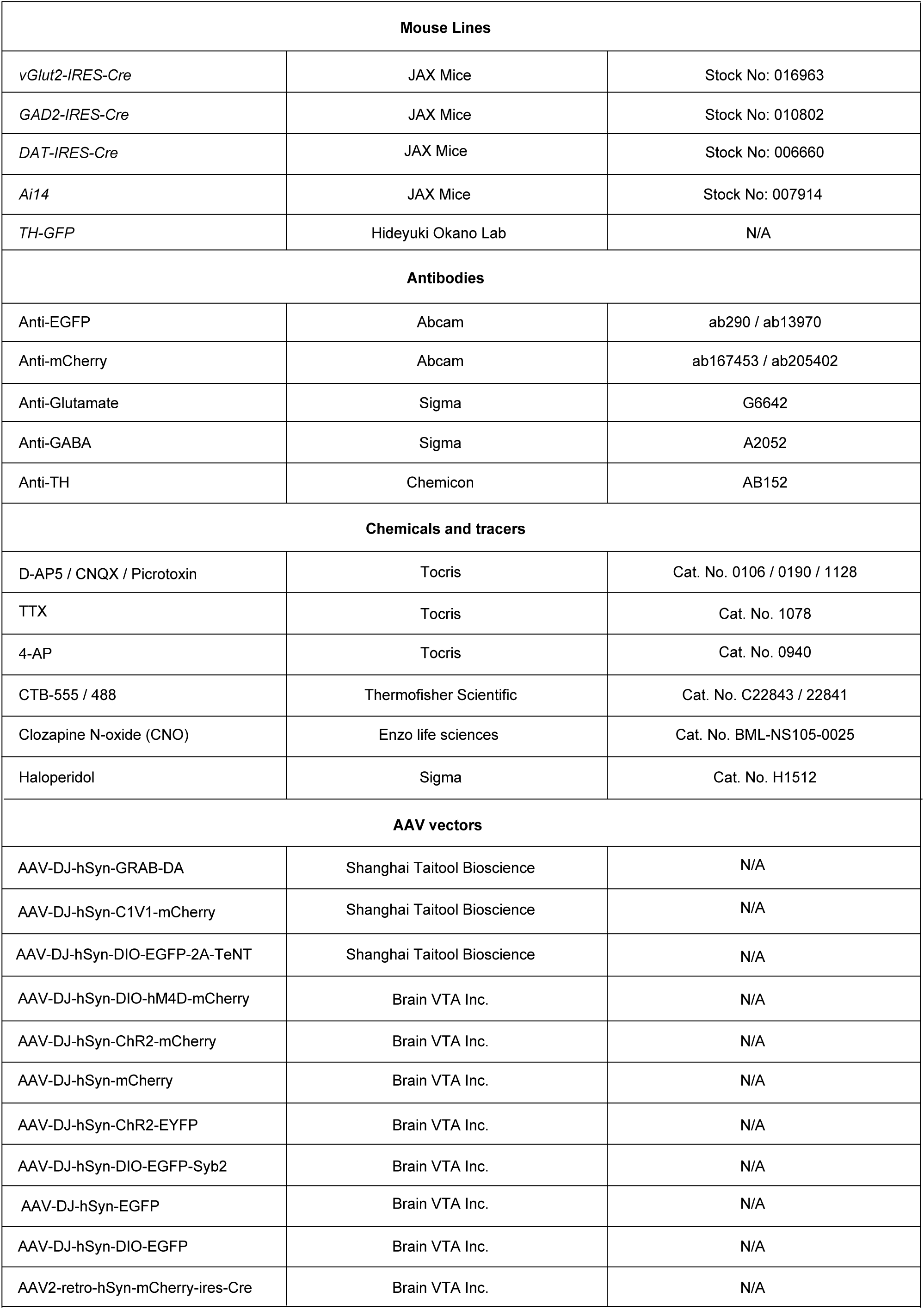
Mouse lines and reagents.

**Table S2.** Summary of all experimental designs.

| Figures | Aims | Mouse lines | AAV injection & optical fiber implantation | Type of data |
| --- | --- | --- | --- | --- |
| **Figure 1**<br>Figure S1 | Cell-type-specific mapping of SC-SNc pathway | *vGlut2-ires-Cre*<br>*GAD2-ires-Cre*<br>*WT* | *AAV-hSyn-DIO-EGFP-Syb2* injected into the SC<br>*CTB-555* injected into the SNc of WT mice | Histology |
| Figure S2 | Segregation of SC-SNc and SC-VTA pathways | *WT* | *CTB-555* / *CTB488* injected into the SNc and VTA, respectively<br>*CTB-555* / *CTB488* mixture injected into the SNc | Histology |
| Figure S3 | Segregation of SC-SNc and SC-ZI pathways | *WT* | *CTB-555* / *CTB488* injected into the SNc and ZI, respectively | Histology |
| **Figure 2**<br>Figure S4 | SNc-projecting SC neurons encode locomotion speed | *WT* | *AAV-hSyn-ChR2-mCherry* injected into the SC<br>Optical fibers implanted above the SNc | Single-unit recording |
| **Figure 3**<br>Figure S5 | Activation of the SC-SNc pathway promotes appetitive locomotion | *WT* | *AAV-hSyn-ChR2-mCherry* injected into the SC<br>Optical fibers implanted above the SNc | Behavior<br>Histology<br>Slice physiology |
| **Figure 4**<br>Figure S6 | The SC-SNc pathway is selectively required for appetitive locomotion | *WT* | *AAV2-retro-mCherry-IRES-Cre* injected into the SNc<br>*AAV-hSyn-DIO-EGFP-2A-TeNT* injected into the SC<br>*AAV-hSyn-DIO-EGFP* (Ctrl) injected into the SC | Behavior<br>Histology |
| **Figure 5**<br>Figure S7 | Genetic labeling of different cell types in the SNc<br>Synaptic innervation on SNc dopamine and non-dopamine neurons | *vGlut2-IRES-Cre*<br>*GAD2-IRES-Cre*<br>*Ai14*<br>*TH-GFP* | *AAV-hSyn-ChR2-mCherry* injected into the SC | Histology<br>Slice physiology |
| **Figure 6** | Activation of the SC-SNc pathway trigger dopamine release in the dorsal striatum | *WT* | *AAV-hSyn-C1V1-mCherry* injected into the SC<br>*AAV-hSyn-GRAB-DA* injected into the dorsal striatum<br>Optical fibers implanted above the SNc and dorsal striatum | Fiber photometry<br>Histology<br>Slice physiology |
| **Figure 7**<br>Figure S8 | SNc dopamine neurons mediates the effect of the SC-SNc pathway activation | *DAT-IRES-Cre* | *AAV-hSyn-ChR2-EYFP* injected into the SC<br>*AAV-hSyn-DIO-hM4D-mCherry* injected into the SNc<br>*AAV-hSyn-DIO-mCherry* (Ctrl) injected into the SNc<br>Optical fibers implanted above the SNc | Behavior<br>Histology<br>Slice physiology |

**Table S3.**
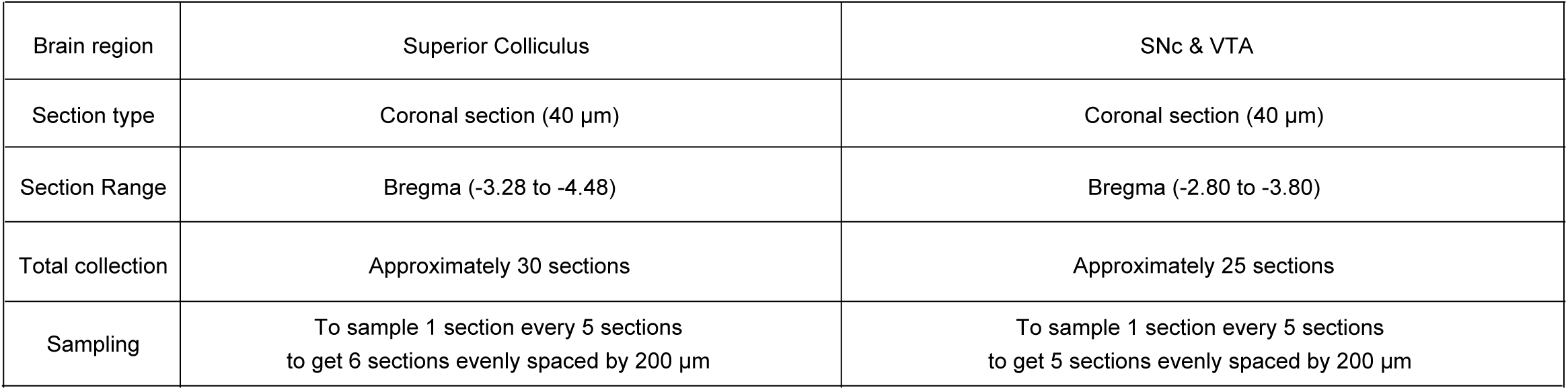
Summary of cell counting strategy.

|  |  |  |
| --- | --- | --- |
| Brain region | Superior Colliculus | SNc & VTA |
| Section type | Coronal section (40 µm) | Coronal section (40 µm) |
| Section Range | Bregma (-3.28 to -4.48) | Bregma (-2.80 to -3.80) |
| Total collection | Approximately 30 sections | Approximately 25 sections |
| Sampling | To sample 1 section every 5 sections<br>to get 6 sections evenly spaced by 200 µm | To sample 1 section every 5 sections<br>to get 5 sections evenly spaced by 200 µm |

**Table S4.**
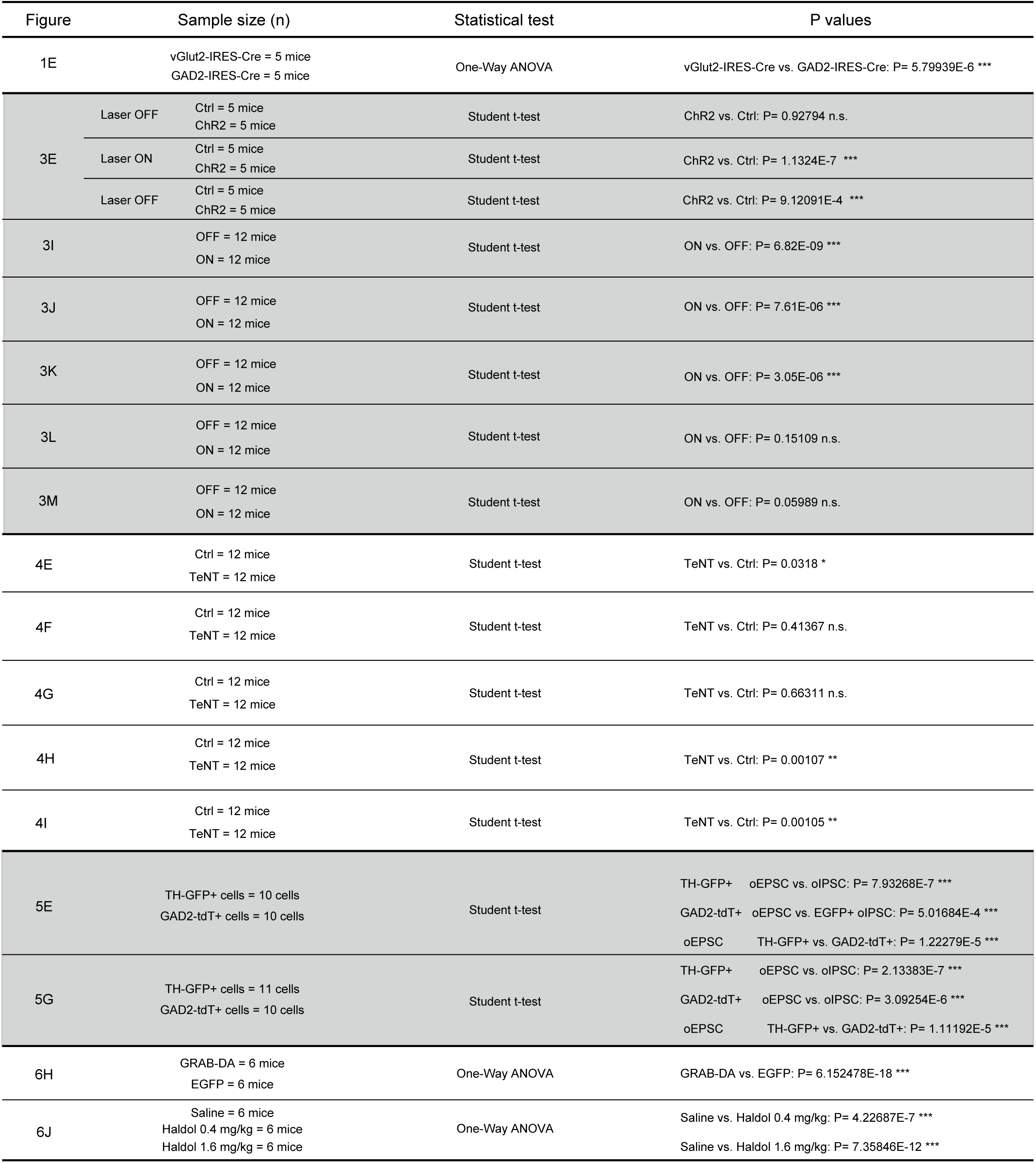
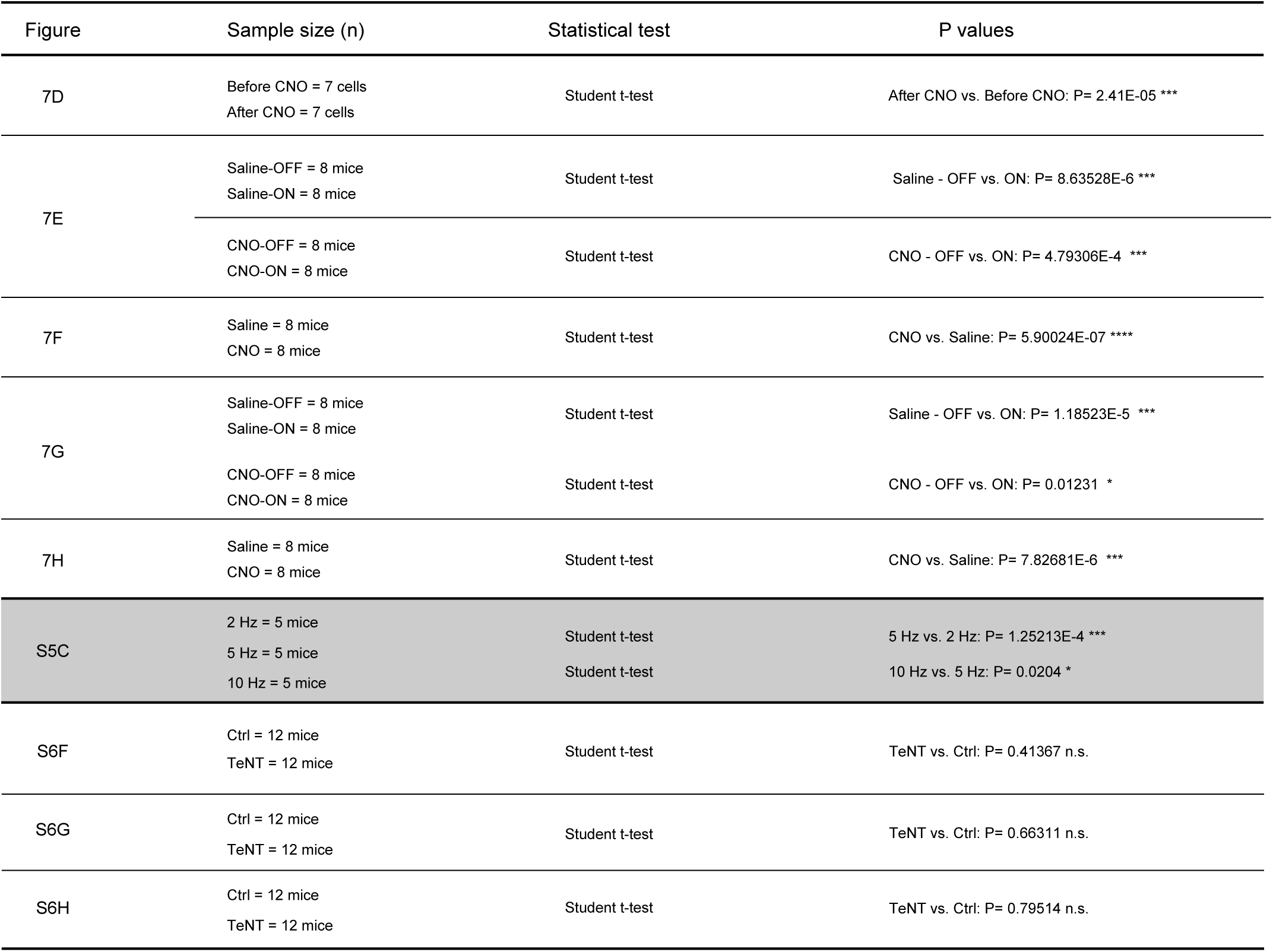
Summary of statistical analyses.

